# Comparative Features of Calretinin, Calbindin and Parvalbumin Expressing Interneurons in Mouse and Monkey Primary Visual and Frontal Cortices

**DOI:** 10.1101/2023.02.27.530269

**Authors:** Maria Medalla, Bingxin Mo, Rakin Nasar, Yuxin Zhou, Junwoo Park, Jennifer I Luebke

## Abstract

Much is known about differences in pyramidal cells across cortical areas and species, but studies of interneurons have focused on comparisons within single cortical areas and/or species. Here we quantified the distribution and somato-dendritic morphology of interneurons expressing one or more of the calcium binding proteins (CaBPs) calretinin (CR), calbindin (CB) and/or parvalbumin (PV) in mouse (*Mus musculus*) versus rhesus monkey (*Macaca mulatta*) in two functionally and cytoarchitectonically distinct regions- the primary visual and frontal cortical areas. The density, laminar distribution and morphology of interneurons were assessed in serial brain sections using immunofluorescent multi-labeling, stereological counting and 3D reconstructions. There were significantly higher densities of CB+ and PV+ neurons in visual compared to frontal areas in both species. The main species difference was the significantly greater density and proportion of CR+ interneurons and lower extent of CaBP co-expression in monkey compared to mouse cortices. Cluster analyses revealed that the somato-dendritic morphology of layer 2-3 inhibitory interneurons is more dependent on CaBP expression than on species and area. Only modest effects of species were observed for CB+ and PV+ interneuron morphologies, while CR+ neurons showed no difference. By contrast to pyramidal cells which show highly distinctive area- and species-specific features, here we found more subtle differences in the distribution and features of interneurons across areas and species. These data yield insight into how nuanced differences in the population organization and properties of neurons may underlie specializations in cortical regions to confer species and area-specific functional capacities.

**Key Points:**

- Somato-dendritic morphology of distinct interneurons did not substantially scale and vary across areas and species- differences were mainly dependent on CaBP expression.
- Cortical diversity in inhibitory function across areas and species is thus likely to be derived from differential laminar distribution and densities of distinct interneuron subclasses.
- In contrast to pyramidal cells which differ widely in distribution and morphology across areas and species, the features of interneurons appears to be relatively more conserved across areas and species.

## INTRODUCTION

Neocortical circuits are comprised of interconnected networks of diverse glutamatergic excitatory neurons and GABAergic inhibitory interneurons. The seminal work of Pandya and other investigators established that the composition and configuration of these networks in different cortical areas are reflected in their distinctive cytoarchitectonic and connectional features (Barbas & Pandya, 1989; O’Kusky & Colonnier, 1982; Petrides et al., 2012; Rockland & Pandya, 1979; Van De Werd et al., 2010; Wang & Burkhalter, 2007) (review: Pandya, 1985; Yeterian et al., 2012). While representing only 20-30% of all cortical neurons, GABAergic inhibitory interneurons play a crucial role in shaping the activity of these networks that mediate area- and species-specific cortical functions (DeFelipe et al., 2013; Isaacson & Scanziani, 2011; Kubota, 2014; Lourenco et al., 2020; Markram et al., 2004). It is well established that the dendritic arbor size and number of dendritic spines of pyramidal cells differ markedly in specific cortical areas of primates vs. rodents, with these differences being particularly evident in frontal cortical areas (review: Luebke, 2017). In frontal association cortices, pyramidal cells increase in size or “scale” by several orders of magnitude from the rodent to the macaque (Ballesteros-Yanez et al., 2007; Elston et al., 2011; Elston & DeFelipe, 2002; Gilman et al., 2017) (reviews:Elston et al., 2011; Luebke, 2017; Wittenberg, 2012). Further, our group has recently shown that in rhesus monkeys (Gilman et al., 2017), morpho-electric properties of pyramidal neurons differ markedly between the lateral prefrontal cortex (LPFC), a higher-order association cortex, and the primary visual cortex (V1), a primary sensory area. In contrast, in rodents, pyramidal neurons in frontal and visual cortices (FC and V1) are largely homogeneous with respect to dendritic topology and biophysical properties (Gilman et al., 2017). These fundamental differences in regionally specific properties of rodent and primate pyramidal cells highlight the implausibility of extrapolation from mouse to primate neurons and cortical networks. Far less is known about comparative morphological features of GABAergic inhibitory interneurons between rodents and primates, especially across distinct cortical areas.

Different types of inhibitory interneurons are characterized by distinctive electrophysiological, neurochemical and structural features, and their functional roles within neocortical circuits are largely determined by their specific axonal distributions and postsynaptic targets (review:Cauli et al., 1997; DeFelipe et al., 2013; Kawaguchi & Kubota, 1997; Kubota et al., 2016; Tremblay et al., 2016). Functionally and morpho-electrically distinct interneuron types are found in all layers of the neocortex, where they exert both powerful and nuanced inhibitory control over neighboring excitatory neurons and on other inhibitory neurons. The distribution and detailed features of cortical interneurons have been characterized by taking advantage of the presence of largely non-overlapping specific molecular markers present in the different types of GABAergic interneurons. Importantly however, markers characteristic of diverse non-overlapping interneuron populations differ between rodents and primates. In the mouse and rat cortex, cortical interneurons can be classified by the presence of calcium-binding proteins parvalbumin (PV) or calretinin (CR), by 5HT3A receptor expression or by expression of neuropeptides such as somatostatin (SOM), vasoactive intestinal peptide (VIP) or neuropeptide Y (NPY) (review: Keller et al., 2018). Cortical GABAergic interneurons in primates such as the rhesus macaque and human do not broadly express neuropeptides but are differentiated by expression of calcium-binding proteins (CaBPs) PV, CR or Calbindin-D28K (CB) (Conde et al., 1994; DeFelipe; Disney & Aoki, 2008; Dombrowski et al., 2001; van Brederode et al., 1990; Zaitsev et al., 2005; Zaitsev et al., 2009). The characteristic morphological features of these neurochemically distinct interneurons are also distinctive. In both rodents and primates, CR+ neurons are Cajal-Retzius cells (located in layer 1 during development), small double bouquet cells or, most commonly, bipolar cells which target the dendritic shafts and spines of pyramidal cells (Dzaja et al., 2014). CR+ interneurons in monkeys are thought to be analogous to VIP+ neurons in rodents, which mainly target other inhibitory interneurons thereby exerting a disinhibitory effect on pyramidal cells (reviews: DeFelipe, 1997; Dzaja et al., 2014; Meskenaite, 1997). CB+ neurons are Martinotti cells, which principally target the apical tufts of pyramidal cells, bipolar cells, and neurogliaform cells which target pyramidal cell dendrites and spines (reviews: DeFelipe, 1997; DeFelipe et al., 2013). CB+ neurons in monkey are analogous to SOM+ neurons in rodents (Barinka et al., 2012; Gonchar et al., 2007). Finally, PV+ neurons are either multipolar basket cells that target the somata of pyramidal cells or Chandelier (axo-axonic) cells that target the axon initial segment of pyramidal cells. Of note, some pyramidal neurons express low levels of these calcium binding proteins as well -for example, layer 3 pyramidal cells express CB in multiple species- but these are readily discriminable from the morphologically distinct classes of interneurons (Cauli et al., 2014; Conde et al., 1994; DeFelipe et al., 1990; Hof et al., 1999). Different interneuron subtypes exhibit distinctive distributions across the six neocortical layers- CR+ and CB+ interneurons neurons are primarily located superficially in layers 1-3 while PV+ neurons are localized principally in layers 2-4. (Conde et al., 1994; DeFelipe; Disney & Aoki, 2008; Dombrowski et al., 2001; Kooijmans et al., 2014; Kooijmans et al., 2020; Tremblay et al., 2016; van Brederode et al., 1990; Zaitsev et al., 2005; Zaitsev et al., 2009). How these neurochemically distinct subpopulations of interneurons compare across species and cortical areas is poorly understood.

There is a higher proportion of GABAergic neurons in the primate neocortex (∼25-34% of all neurons) compared to the rodent neocortex (∼15-25% of all neurons) (Beaulieu, 1993; DeFelipe, 2002; Gabbott & Bacon, 1996b; Gabbott et al., 1997; Gonchar & Burkhalter, 1997; Meinecke & Peters, 1987). This difference is likely due to a significantly higher proportion CR+ interneurons in layers 2 and 3 in the primate (12%) compared to rodent (4%) (Gabbott & Bacon, 1996a, 1996b; Gabbott et al., 1997) (review:Dzaja et al., 2014). On the other hand, PV+ interneurons have been reported to comprise a lower proportion of all interneurons in the primate compared to rodent neocortex (Conde et al., 1994; Desgent et al., 2005; Gabbott & Bacon, 1996b; Gabbott et al., 1997; Glezer et al., 1998; Gonchar & Burkhalter; Kawaguchi & Kubota, 1997). Very few studies have directly quantitatively compared the distribution of all 3 subtypes of CaBP positive interneurons in the same area of rodent and primate cortices in the same study. In one of the only such studies, Kooijmans and colleagues (Kooijmans et al., 2020) compared the distribution, relative proportion and size of CR+, CB+ and PV+ interneurons and of interneurons in which these markers were colocalized in V1 of the mouse and the rhesus monkey. This study reported a significantly higher proportion of PV+ and of CB+ neurons and a lower proportion of CR+ neurons in monkey V1 compared to mouse. In addition, fewer neurons were positive for multiple calcium-binding proteins and a more spatially segregated (to layers 2-4) distribution of PV+ and CB+ neurons was seen in monkey compared to mouse V1. (Kooijmans et al., 2020) confirmed previous reports of a higher number of CaBP positive inhibitory interneurons in primates compared to mice (reviews: Defelipe, 2011; Dzaja et al., 2014). Comparative studies of the morphological features of these CaBP expressing interneurons is also limited but indicate that areal or species differences are much more subtle than those seen in pyramidal cells. For example, several studies report no significant between-area and between-species differences in the size of CR+ neurons (Gabbott & Bacon, 1996a; Gabbott et al., 1997; Kooijmans et al., 2020) or PV+ neurons (Disney & Reynolds, 2014; Povysheva et al., 2008). Other studies have report small but significant between-species and between-area differences in visual cortices, with PV+ interneuron cell bodies slightly larger in monkey than in mouse V1 (Kooijmans et al., 2020) and in monkey area V2 compared to V1 (Sherwood et al., 2007).

While a number of previous studies have characterized the distribution of interneurons in specific brain areas in mice or in monkey, these have typically been limited to a single brain area such as V1 (e.g. Kooijmans et al., 2020) or across different regions in a single species (e.g. Dombrowski et al., 2001). What remains unclear is whether the distribution and morphology of CR+, CB+ and/or PV+ neurons differ by brain region within and between species (mouse and monkey). Here we extended previous studies by comparing interneurons using the same protein markers in two functionally distinct brain areas, V1 and FC in mouse and V1 and LPFC in rhesus monkey. We compared 1) the total density and laminar localization of neurons containing the calcium binding proteins CR, CB and PV; 2) the density and distribution of neurons co-expressing two or more of these CaBPs, and 3) the comparative size and primary dendritic branch number, polarity and orientation of these neurons. The degree to which individual and populations of mouse and monkey interneurons are similar or differ has major implications for the generalizability of ‘canonical’ cortical circuits across rodent to primate species and distinct cortical areas.

## MATERIALS & METHODS

### Subjects

Subjects included 3 adult C57BL/6J mice (*Mus musculis*; all males; 8-9 months of age) and 3 adult rhesus monkeys (*Macaca mulatta*; 1 male, 2 female; 17-20 years of age), which were part of larger ongoing parallel studies. Mice were acquired from private vendors. Monkeys with complete health records were acquired from national primate research facilities and from private vendors and were part of our larger studies of normal brain aging. All procedures were approved by the Boston University Institutional Animal Care and Use Committee and were conducted in strict accordance with the guidelines established by the NIH Guide for the Care and Use of Laboratory Animals.

Whole brain tissue was fixed and extracted from mice as described previously (LeBlang et al., 2020). Mice were sedated with sodium pentobarbital (6 mg/kg, to effect), and perfused intracardially with 4% paraformaldehyde (PFA, 36°C). After removal of the dorsal skull cap, the brain was post-fixed *in situ*, in 4% PFA overnight at 4°C. The brain was then extracted from the skull and post-fixed in 4% PFA for an additional 48 hours, at 4 °C. The brains were then serially sectioned in the coronal plane at 100 μm using a Leica VT1000S Vibratome. Sections used in the study included area Fr2 from the frontal region and area V1 from the occipital region.

Whole brain tissue was fixed and extracted from monkeys using our 2-stage perfusion procedure described previously (Medalla et al., 2021; Medalla et al., 2017). Briefly, monkeys were sedated with ketamine hydrochloride (10 mg/kg) and subsequently anesthetized with sodium pentobarbital (15 mg/kg, to effect). Then, monkeys were perfused through the ascending aorta with ice-cold Krebs–Henseleit buffer, which contained, in mM: 6.4 Na_2_HPO_4_, 1.4 Na_2_PO_4_, 137 NaCl, 2.7 KCl, 5 glucose, 0.3 CaCl_2_, and 1 MgCl_2_, pH 7.4 (Sigma-Aldrich). Brain tissue blocks about 10 mm^3^ in size were taken from both LPFC (ventral area 46) and lateral V1 and were immersion-fixed immediately in 4% PFA for over 48 hours at 4C. Fixed tissue blocks were then sectioned in the coronal plane at 100 μm, in ice-cold 0.1M PB (pH 7.4). Serial brain sections from both mice and monkeys were stored in antifreeze solution, consisting of 30% ethylene glycol, 30% glycerol, 40% 0.05 M Phosphate Buffer (PB), at −20°C, until use.

### Regions of Interest

Regions of Interest (ROIs) included functionally analogous frontal and visual areas in mice and monkeys based on our previous studies (Gilman et al., 2017; Hsu et al., 2017), and cytoarchitectonic maps for monkey (Barbas & Pandya, 1989; O’Kusky & Colonnier, 1982; Pandya, 1985; Petrides et al., 2012; Rockland & Pandya, 1979) and mouse cortices (Anastasiades et al., 2019; Van De Werd et al., 2010; Wang & Burkhalter, 2007; Xu et al., 2016) (Allen Brain mouse reference atlas; atlas.brain-map.org) (see Fig. 1a). Rhesus monkey frontal ROI was in Brodmann area 46 located in the ventral bank of the principal sulcus within the LPFC (Barbas & Pandya, 1989), with blocks sampled within the caudal half of the principal sulcus (Fig. 1a). How and whether frontal areas in rodents correspond to monkey prefrontal cortical areas has been a long-standing debate in the field. In particular, it is questionable whether there is a mouse brain region homologous to the primate LPFC (Uylings et al., 2003). While recognizing this caveat, here and in our previous work, we assessed the dorsolateral FC area Fr2 of the mouse (atlas.brain-map.org) (Gilman et al., 2017), which is within an anatomical location analogous to the monkey LPFC. For primary visual cortex, the rhesus monkey primary visual cortex ROI was within the lateral region of V1, Brodmann area 17 (Rockland & Pandya, 1979) (Fig. 1a). The mouse V1 ROI was identified based on Allen Brain mouse reference atlas (atlas.brain-map.org), as in our previous studies (Gilman et al., 2017; Hsu et al., 2017) (Fig. 1a).

**Figure 1.**
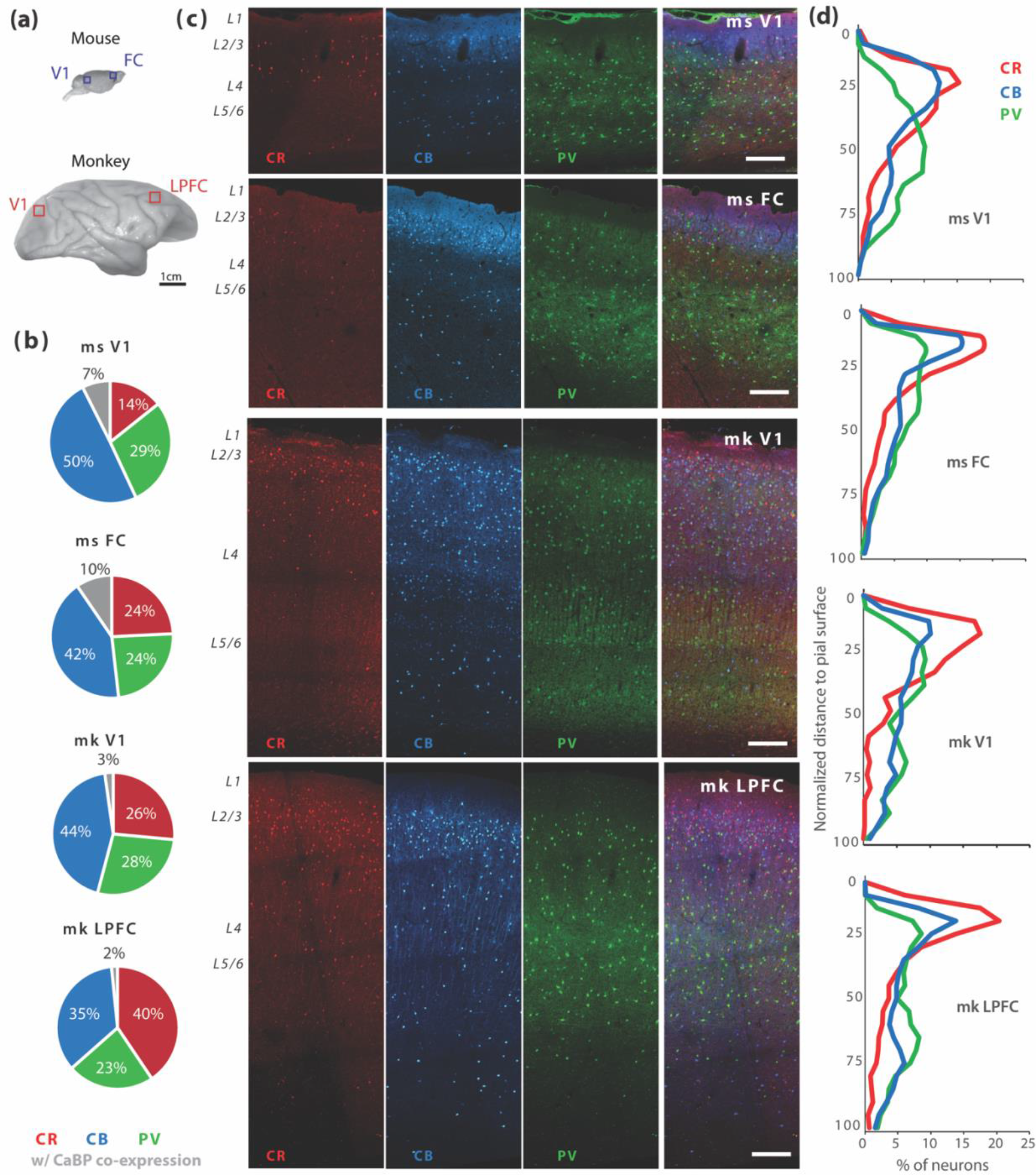
**Distribution of CR+, CB+ and PV+ interneurons in mouse and monkey visual and frontal cortices**. **a)** Lateral surface images of mouse and monkey brains showing locations of sampled visual and frontal cortical areas. **b)** Pie charts showing proportions of interneurons expressing CR, CB, PV or a combination of two or more CaBPs. **c)** Confocal image montages throughout the cortical depth from pia to white matter of mouse and monkey frontal and visual cortices, showing individual and merged channels (far right panels) of CR, CB and PV triple labeling. Scale bars: 200µm. **d)** Cortical depth distribution profiles of CR+, CB+, and PV+ interneurons across normalized distance from pia, in mouse and monkey frontal and visual cortices.

### Immunohistochemistry

Triple labeling immunohistochemical (IHC) fluorescence labeling experiments were conducted to assess the density, location and morphology of interneurons positive for the calcium binding proteins CR, CB, and/or PV. For each animal and area, 3 sections ∼500um apart were selected for IHC experiments. Sections were first washed in 0.01M phosphate-buffered saline (PBS, 3x10 minutes, 20°C) and then incubated in 50mM glycine in 0.01M PBS at 20°C for 60 minutes. Antigen retrieval was performed by incubation of sections in 10mM Sodium Citrate Buffer (pH ∼8.5) at 60-70°C for 20 minutes, followed by 30 minutes cool-down to ∼20°C. After rinsing (3x10 minutes in PBS), sections were incubated in pre-block solution [(0.01M PBS, 5% Bovine Serum Albumin (powder), 5% Normal Goat Serum; 0.2% Triton X-100] for 60 minutes at 20°C. Sections were then incubated in a combination of primary antibodies diluted in the diluent buffer containing 0.1M PB; 0.2% Bovine Serum Albumin-acetylated; 1% Normal Goat Serum; 0.1% Triton X-100. Two sets of batch-processing experiments were conducted with the following primary antibody combinations. In the first set of experiments, sections were co-incubated in primary antibodies against CR (rabbit anti-CR/Calbindin D-22k, 1:1000; Swant, 7697; RRID AB_2721226; Swant Inc, Burgdorf, Switzerland) and PV (mouse anti-PV, 1:1000; Swant 235, RRID AB_10000343). In a second set of experiments designed to include CB, and also to allow for CaBP colocalization quantification, sections were co-incubated in rabbit anti- CB (Calbindin D-28k, 1:2000; Swant #CB38; RRID AB_10000340) together with a mouse anti-PV (1:1000; Swant 235, RRID AB_10000343) and a new guinea pig anti-CR (Calbindin D-22k, 1:500; Swant CRgp7). For each combination of primary antibodies, incubation of sections was done via 2x 10-minute microwave sessions at 150W, 40°C using the PELCO Biowave (Ted Pella, Inc., Redding, CA, USA) followed by a 4°C overnight incubation for 72 hours. Sections were then washed in 0.01M PBS (3x10 minutes), then incubated in secondary antibody solution at 4°C for 24 hours. The following combination of secondary antibodies conjugated to distinct Alexa fluor fluorescent probes were used: Alexa 488 conjugated donkey anti-guinea pig IgG (1:200), Alexa 546 conjugated donkey anti-mouse IgG (1:200), Alexa 647 donkey anti-rabbit IgG (1:200). After a final rinse in 0.1M PB (3x10 minutes 4°C), sections were mounted on glass slides and cover slipped with ProLong Gold Antifade (Thermo Fisher Scientific, Waltham, MA, USA). Control experiments omitting the primary antibodies but incubating only in the combination of secondary antibodies in each set of tissue were performed and yielded no labeling.

The antibodies used have been validated by the manufacturer and are widely used in the literature. The rabbit polyclonal CB antibody (Swant #CB38; RRID AB_10000340) was raised against recombinant rat calbindin D-28k, and recognizes a single band of approximately 27 -28 kDa in immunoblots (see manufacturer’s datasheet). The PV antibody is a mouse monoclonal antibody (Swant 235, RRID AB_10000343) produced via hybridization of mouse myeloma cells with spleen cells from mice immunized with parvalbumin purified from carp muscles (see manufacturer’s datasheet). The rabbit CR antibody (Swant, 7697; RRID AB_2721226) was raised against recombinant human calretinin (calbindin D-22k) containing a 6-his tag at the N-terminal. The guinea pig CR antibody (Swant, CRgp7) was raised against recombinant mouse calretinin (calbindin D-22k). Both CR antibodies have been validated to react with monkeys, and specificity has been tested using immunoblotting methods and using tissue from CR knock-out mice (see manufacturer’s datasheet).

The rabbit CB, mouse PV, and rabbit CR antibodies have been well documented to work in rhesus monkeys and rodents in the published literature (e.g. Barbas et al., 2005; Conde et al., 1994; DeFelipe et al., 1999; Dombrowski et al., 2001; Kondo et al., 1994; Kondo et al., 1999; Kooijmans et al., 2020; Povysheva et al., 2008; Zaitsev et al., 2009). Although the CR guinea pig antibody (Swant, CRgp7) has been validated by the manufacturer to react in monkey tissue, it has not been widely used in primates in the literature. Thus, we performed a control validation dual labeling experiment using guinea pig anti-CR (1:500, Swant, CRgp7) together with rabbit anti-CR (1:1000; Swant, 7697; RRID AB_2721226) in the same tissues from one monkey and one mouse case, using the identical protocol as for the actual experiments (see above). For each of the areas examined except for monkey V1, the majority (86-91%) of the cells were dual labeled with CR-rabbit and CR-guinea pig. In monkey V1 however only 50% of the cells were dual labeled with CR-rabbit and CR-guinea pig (Supplementary Material Fig. S1). We noted that the CR-guinea pig antibody consistently stains a smaller population of neurons compared to CR-rabbit, and this effect is larger in monkey V1. Thus, we used two sets of immunolabeling experiments – the dual labeled and triple labeling experiments, which used CR-rabbit and CR-guinea pig respectively – to quantify CR+ densities and colocalization between proteins.

### Confocal Laser Scanning Microscopy

Brain sections were imaged using a LEICA SPE laser-scanning confocal microscope (Lieca Biosystems, Wetzlar, Germany). For counting the density of cell bodies single-labeled or co-labeled with either CR, CB and/or PV image stacks were obtained using a 20x 0.5 N.A water immersion objective. A tile scan was performed to capture a 400µm wide cortical column from the pia to the white matter for each area and section (Fig. 1). Image stacks were acquired at 0.538 x 0.538 µm pixel size and 1 or 2 µm z-step size. Three channels were scanned separately using three diode lasers with the following wavelengths: 488 nm for green (Alexa 488), 561 nm for red (Alexa 546), and 647nm for far-red (Alexa 647) labels. After imaging, image stacks were deconvolved using the AutoQuant software as described (Gilman et al., 2017; Tsolias & Medalla, 2022). For each case, 3 serial sections were imaged through each ROI. Thus, a total of 18 (3 sections x 2 areas x 3 animal cases) sections of mouse brain tissue and 18 (3 sections x 2 areas x 3 animal cases) sections of monkey brain tissue was analyzed in this study.

### Quantification of the distribution of single-labeled and co-labeled neurons

Neuronal counts across the entire cortical depth from pia to white matter of each image z-stack montage were obtained using Neurolucida 360 (NL360) version 2019.1.1 (MBF Bioscience, Williston, VT, USA; Fig. 1). A total of 34425 neurons were counted in mouse V1 (n cells = 9273), mouse FC (n = 13865), monkey V1 (n = 11287) and monkey LPFC (n = 9273) from 3 cases in each species. All neurons were exhaustively counted in each z-stack using 3D stereological counting rules, by placing exclusion and inclusion planes at the top and bottom each z-stack to avoid counting errors due to cell plucking and slicing during sectioning, as described (Tsolias & Medalla, 2022). We marked all strongly immunolabeled neurons with non-pyramidal morphology within the entire z-stack. Colocalization was determined by the overlap of two- or three- color channels within an identified cell body in the same z-plane. Immunolabeled neurons were marked as follows: CR+, CB+, PV+ (single-labeled neurons), CR+CB+, CR+PV+ or CB+PV+ (dual-labeled neurons). Since the type of CR antibody showed some methodological variability, with CR-guinea pig antibody labeling fewer cells than the CR-rabbit antibody, we counted total densities of CR using the experiment that used the CR rabbit antibody (Supplementary Material Fig. S1). Density of colocalization was based on both the triple labeling experiments that used the CR guinea pig antibody, and dual labeling experiments using the CR rabbit antibody, which yielded consistent results.

To assess the cortical depth distribution profile of each immunolabeled neuronal population, open contours to delineate the pial surface and white matter (WM) were traced for each section. The location of each marker within the counting volume and its distance from the pia was measured. The cortical depth of each marker, expressed as a normalized (%) distance from pia, was calculated by dividing the distance from pia of each marker to the total cortical depth (distance between pia and WM contours). A histogram of normalized neuronal counts as a function of % distance from pia was obtained for each individual case, depicting the cortical depth profile of each neuronal population.

For each population, the total density and the density in upper layers 1-3 versus deeper layers 4-6, were assessed. Closed contours to delineate these laminar ROIs were marked on each section, guided by depth measurements obtained from matched DAPI (4′,6-diamidino-2- phenylindole) labeled sections. The distance of each layer from the pia in each area was measured from DAPI reference sections and published data (Anastasiades et al., 2019; Xu et al., 2016) (Table 1). Specifically, layer boundaries for monkey PFC and V1 were measured from adjacent DAPI labeled sections. The layer boundaries for mouse V1 and FC were adopted from Xu et al. 2016 and Anastasiades et al., 2018, respectively (Table 1). The total volume of the sampled column and the volume of each laminar ROI were calculated (total surface area of closed contours x number of optical slices x z step size). For each subject and area, densities were calculated by pooling total neuronal counts from 3 sections and dividing this by the total ROI volume from the 3 sections.

**Table 1:** Boundaries of layers 1-6 for species and areas, based on percent distance from pia.

|  | Monkey LPFC | Monkey V1 | Mouse FC<br>(Anastasiades et al.,<br>2019) | Mouse V1<br>(Xu et al., 2016) |
| --- | --- | --- | --- | --- |
| Layer 1 | 0%-7% | 0%-6% | 0%-15% | 0%-12.5% |
| Layer 2-3 | 7%-40% | 6%-36% | 15%-40% | 12.5%-35% |
| Layer 4 | 40%-50% | 36%-68% | 40%-50% | 35%-40% |
| Layer 5 | 50%-71% | 68%-79% | 50%-75% | 40%-65% |
| Layer 6 | 71%-100% | 79%-100% | 75%-100% | 65%-100% |

### 3D analyses of somato-dendritic morphology and dendritic arbor laminar span

Immunolabeled neurons with complete somata and substantial labeling of processes within the image z-stack were identified in L2-3 for 3D reconstruction. Cell inclusion criteria for 3D reconstruction were as follows: soma complete in the z axis, at least 50% of the primary dendrites extend ∼2x the diameter of the cell body, and at least 2 dendritic branches spanning vertically and 2 spanning horizontally were complete within the z-stack.

A total of 386 cells were reconstructed (an average of 30 cells per area and CaBP+ cell type), with soma-to-pia distance of cells that were within the boundaries of L2-3 as follows: Cells within 250-500 µm deep from the pia for monkey V1, 200-700 µm deep for monkey LPFC, and 200-400 µm deep for mouse V1 and FC (Gilman et al., 2017; Medalla & Barbas; Van De Werd et al., 2010). The somata and visible dendrites of neurons were reconstructed in 3D using Neurolucida 360 (NL360) version 2019.1.1 (MBF Bioscience) as described in our previous work (Guillamon-Vivancos et al., 2019; Medalla et al., 2021). All somata were automatically detected and dendrites were manually traced throughout the z-stack. Exporting and analyses of the 3D tracing data was done in Neurolucida Explorer (MBF Bioscience). For each neuron the soma volume, number of primary dendrites and their polarity, and the maximum vertical and horizontal dendritic span were quantified. The vertical span was measured by drawing a straight line perpendicular to the pial surface that spanned the most superficial and deepest tips of the dendritic arbor. The horizontal span was measured by drawing a straight line parallel to the pial surface that spanned the farthest tips of the dendritic arbors. Laminar boundaries delineating L1, L2, L3 and L4 were placed to assess for each cell (with a soma in L2-3) the number of layers spanned by the dendritic arbor.

Dendritic polarity was characterized by first obtaining a radial histogram of all dendritic lengths for each cell in Neurolucida Explorer. This histogram was exported and then a vector component was calculated as a measure of a bias of each cell towards an angular orientation, as described in (Guillamon-Vivancos et al., 2019). This vector quantification was obtained for each cell as follows: first, dendritic length in each angular bin was normalized to the total dendritic length. The resulting value per bin was then multiplied by cos(*θ*) to get the vector x component, and sin(*θ*) to get the vector y component, where *θ* is the midpoint angle of each bin (i.e., for bin = 0 to 10 degrees, *θ* = 5). Then the sum of all x and y components was calculated as a measure of polarity.

### Statistics

All data were exported to Microsoft Excel spreadsheets and analyzed using Excel and MATLAB (MathWorks, Natick, MA, USA). The following neuronal counting outcome measures were compared between-areas, species, CaBP and layers: the normalized number of counted neurons as a function of normalized distance of each marker from pia; density in the total ROI volume, and density by laminar compartment, superficial (1-3) versus deep (4-6) layers. The following somato-dendritic morphological outcome measures for individual immunolabeled L2 and L3 neurons were compared between-areas, species and CaBP as follows: soma size, number of primary dendrites, number of layers spanned, maximum vertical and horizontal span and polarity.

For neuronal distribution and density measures, data from each section were pooled per area and case, and then the average, standard deviation, standard error, and outliers were calculated for the following four groups: mouse V1, mouse FC, monkey V1, and monkey LPFC. For density measures, comparisons were made between cases. For cortical depth measurements (% distance from pia) and somato-dendritic properties of individual neurons, a repeated measures design was used to compare cortical depth bins and morphology of individual neurons.

All statistical analyses were conducted using MATLAB. The Kolmogorov-Smirnov test for normality was employed for each outcome measure. For all measures that have a normal distribution, multiple-comparisons one-way, two-way or three-way ANOVA with Fisher’s LSD post-hoc test was conducted (at p < 0.05 significance level) depending on the nature of the independent and dependent variables tested and compared. One-way ANOVA with multiple comparisons was used to compare the frequency of neurons within each cortical depth bin across the 4 groups by area and species. Two-way ANOVAs were used for independent and interactive between-area and between-species comparisons on the proportions of each CaBP expressing cell subtype (CR+, CB+, PV+ neurons and dual-labeled CRCB, CRPV and CBPV neurons). For the total density of each CaBP expressing cell subtype (CR+, CB+, PV+ neurons), a three-way ANOVA was conducted to compare between-species, between-area and between-CaBP expressing subtype. The laminar (1-3 and 4-6) densities within each subpopulation of interneurons were also compared between-species, between-area and between-layer using a three-way ANOVA. For all ANOVA tests with significant main or interactive effects (*p* < 0.5), Fisher’s LSD multiple comparison post-hoc tests were performed subsequently to specify which pairs were significantly different. Somato-dendritic morphometric properties were compared between individual L2 and L3 cells reconstructed from 2-3 sections/cases per group. These somato-dendritic morphometric features were compared between-area, between-species and between-CaBP cell type using a three-way ANOVA with a repeated measures correction. Chi squared tests within each CaBP subpopulation was used to compare the proportion of cells spanning 1, 2, 3 or 4 layers across the four groups by area and species.

To determine whether these immunolabeled interneuron subpopulations represent distinct morphological subclasses, we analyzed (dis)similarities of interneurons based on the multidimensional dataset of 7 dendritic morphometric outcome measures (soma volume, number of layers spanned by arbors, number of primary dendrites, maximum vertical and horizontal dendritic span, and polarity X and Y projection values) using clustering analyses and dimensional reduction as described (Gilman et al., 2017; Medalla et al., 2020). Each variable was normalized to z-score values, and a distance matrix (based on squared Euclidean distances) was computed. The multidimensional distance matrix was then reduced to two dimensions via non-metric multidimensional scaling (NMDS) which assigns a coordinate value to plot each neuron in in a two-dimensional space. This analysis allows the visualization of multidimensional data in a 2D space, with the distances between each data point representing the similarities in the multidimensional properties—the closer the distance, the more similar the properties of the neurons. We first ran NMDS analyses on the full dataset, with all neurons pooled together, and then ran NMDS plots for each CaBP expressing subpopulation. For each NMDS plot, we then annotated the individual neurons to visualize and see clustering based on categorical grouping variables as follows: 12 subgroups (by area, species and CaBP: CRmsV1, CRmsFC, CRmkV1 CRmkLPFC; CBmsV1, CBmsFC, CBmkV1, CBmkLPFC; PVmsV1, PVmsFC, PVmkV1 PVmkLPFC) separately, by species and area (ms V1, ms FC, mk V1, mk LPFC), by CaBP cell type (CR, CB, PV), by morphological subclass (multipolar vs bipolar), and layer (L2 vs L3). We also visualized values of continuous variables (soma volume, vertical extent, horizontal extent) for each individual neuron on the NMDS plot, using a heatmap gradient scale assigned to the numerical range of each continuous variable.

To validate the bottom-up NMDS clustering scheme, a one-way multivariate analysis of variance (MANOVA) was used for supervised clustering, to compare the relative degree of influence each outcome measure contributes to the separation of the interneuron subgroups by area, species, CaBP, morphology and layer, as described (Medalla et al., 2020). Outcome measures included were as follows: soma volume, number of layers spanned by arbors, number of primary dendrites, maximum vertical and horizontal dendritic span, polarity X and Y projection values. The properties were normalized to z-scores, and two canonical variables—weighted linear sums of all included variables—that best separate out the groups were computed. For each canonical variable, the absolute value of the coefficient represents the degree of influence that each outcome measure contributes to the separation of the groups, with higher absolute values indicating a greater influence.

## RESULTS

### Proportions of CR+, CB+, PV+ interneuron subclasses

We compared the relative proportions of interneurons immunopositive for each of the 3 CaBP in V1 and FC cortical areas in the mouse (ms) and V1 and LPFC in the monkey (mk) (Fig. 1a-c). Fig. 1b pie charts show the relative frequencies of CR+, CB+, PV+ neurons and neurons positive for more than one of these markers in each cortical area. In the mouse, the predominant population was CB+ (50% and 42% in V1 and FC respectively; Fig. 1b top two charts). In the monkey, the predominant population depended on cortical area. In monkey V1 the predominant population was CB+ (44%) while in monkey LPFC, the predominant population was CR+ (40%; Fig. 1b bottom two charts). Two-Way ANOVA (species x area) comparisons revealed significant between-area (main effect, area p = 0.008) and between-species (main effect, species p = 0.0005) differences in the proportion of CR+ neuron subpopulation. Specifically, in monkey V1 compared to LPFC there was a significantly lower proportion of CR+ interneurons (LSD post-hoc, p = 0.012). Between-species comparisons revealed a significantly lower proportion of CR+ interneurons in the mouse compared to monkey independent of cortical area (LSD post-hoc, msV1 vs mkV1 p = 0.013; msFC vs mkLPFC p = 0.0017). Furthermore, interneurons that expressed multiple calcium-binding proteins occurred at a higher frequency in the mouse compared to monkey (main effect species, p = 0.000004; LSD post-hoc, msV1 vs mkV1 p = 0.0002; msFC vs mkLPFC p = 0.00001).

### Cortical depth distribution of CR+, CB+, PV+ interneuron subclasses

We compared the relative depth distribution patterns of CR+, CB+, and PV+ interneuron populations within each area and species (Fig. 1d). Since the pia to white matter distance (cortical depth) differs between areas and between species, we normalized the position of each neuron (distance of the neuron to the pia/total pia to white matter distance; Fig. 1d). Each interneuron subclass exhibited distinctive depth distributions, that were consistent with previous reports (Conde et al., 1994; DeFelipe, 1997; Disney & Aoki, 2008; Dombrowski et al., 2001; Kooijmans et al., 2014; Kooijmans et al., 2020; Tremblay et al., 2016; van Brederode et al., 1990; Zaitsev et al., 2005; Zaitsev et al., 2009); that is, CR+ and CB+ interneurons were distributed superficially- closer to the pia- compared to PV+ interneurons (Fig. 1c, d). One-way ANOVA with multiple comparisons showed that there was a similar distribution of CR+ interneurons across the vertical extent of all 4 areas, with these neurons being highly concentrated within the superficial 20% distance from pia, followed by a sharp decrease (Fig. 1d, red line). In both mouse V1 and FC, CR+ and CB+ neurons showed a prominent peak within the superficial 25% distance from pia, followed by a sharp decrease at around 50% of the pia to white matter distance (Fig. 1c, d). Monkey V1 and LPFC showed a similar superficial distribution of CB+ and CR+ interneurons, however, the superficial peak was less prominent for CB+ neurons in monkey compared to mouse areas (Fig. 1c, d).

There was a significant between-species difference in CB+ interneuron distribution (main effect ANOVA, p=0.006). Mouse V1 showed a higher concentration of CB+ between 15-20% distance from pia compared to monkey V1 (p<0.03, for all LSD post-hoc comparisons), which presented a second peak in CB+ neurons within 80-90% distance from pia (p<0.05; Fig. 1d, blue line). Compared to monkey LPFC, CB+ neurons in mouse FC showed a significantly higher peak at ∼15% distance from pia (LSD post-hoc, p = 0.026), but exhibited a sharp decline, with significantly fewer CB+ neurons between 75-100% distance from pia (p<0.02 for all LSD post-hoc comparisons; Fig. 1d).

Finally, the depth distribution of PV+ interneurons exhibited significant differences between both area and species (main effect ANOVA, p = 0.0021). In both mouse V1 and FC, PV+ neurons showed a unimodal distribution pattern with a wide upper layer peak followed by a gradual decrease between 50-100% of cortical depth. However, this wide peak was more superficial in mouse FC, spanning 25-40% of the cortical depth, but was shifted deeper, between 45-60% in mouse V1 (Fig. 1d, green). By contrast, monkey V1 and LPFC both showed a bimodal PV+ distribution with one superficial peak between 20-40% distance from pia, and a second peak between 70-85% of cortical depth (Fig. 1d, green). Further, between-species comparisons showed an opposite pattern of PV+ depth distribution in mouse versus monkey areas. Mouse FC showed a significantly higher concentration within the superficial 15% of cortical depth compared to monkey LPFC (p<0.02, for all LSD post hoc comparisons). In contrast, monkey LPFC and V1 had significantly greater proportion of PV+ neurons closer to the white matter (between 85-100% of cortical depth), compared to mouse cortices (msV1 vs mkV1, p = 0.002; msFC vs mkLPFC p < 0.002 for all comparisons).

### Total and laminar densities of CR+, CB+, PV+ neurons

We assessed the density (number of neurons/mm^3^) for each interneuron class across the total gray matter depth and within superficial (layers 1-3) and deep (layers 4-6) laminae specifically (Fig. 2). Three-way ANOVA (species*area*CaBP) revealed that significant between-species and between-area differences in total densities of interneuron subpopulations were dependent on CaBP expression (species*CaBP p = 0.0036; area*CaBP p = 0.00035). No between-area differences were found in total CR+ interneuron density within either species. Interestingly however, CR+ density was significantly greater in monkey compared to mouse in both visual and frontal areas (LSD post-hoc, msV1 vs mkV1: CR p = 0.0007; msFC vs mkLPFC: CR p = 0.00008; Fig. 2a). By contrast, significant between-area differences were found in the total density of CB+ and PV+, in both monkey and mouse, but no between-species differences were observed (Fig. 2b, c). In both monkey and mouse, the total density of CB+ neurons was greater in V1 compared to frontal regions (mouse V1 vs FC: CB p = 0.0000099; monkey V1 vs LPFC: CB p = 0. Fig. 2b). Similarly, the density of PV + neurons was greater in V1 compared to FC in mouse (p 0.001; Fig. 2c).

**Figure 2.**
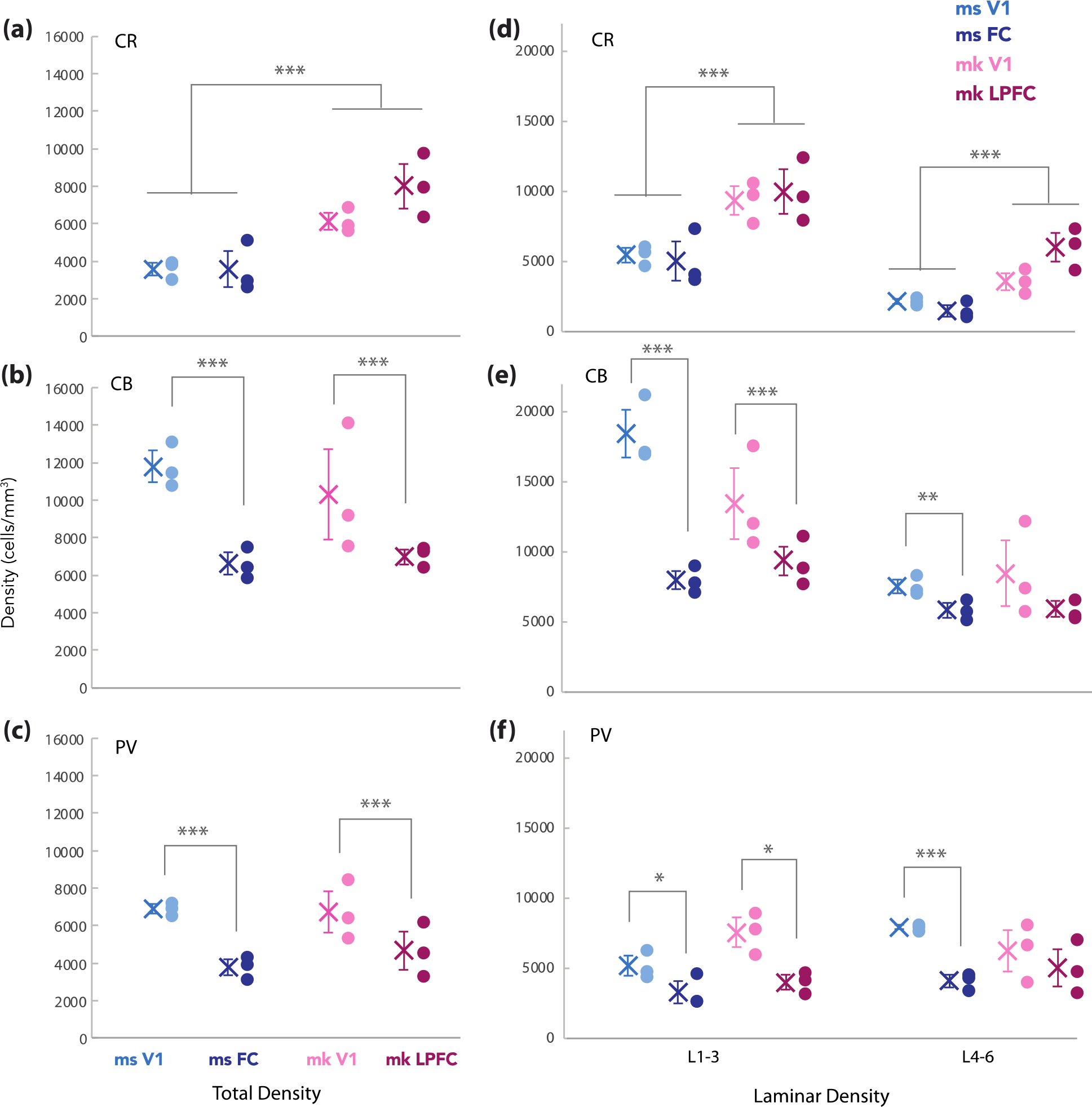
Total and laminar densities of CR+, CB+ and PV+ interneurons. a-c) Total density of neurons immunolabeled for CR, CB and PV and, **d-f)** Density of CR+, CB+, and PV+ interneurons within superficial (1-3) and deep (4-6) layers.

We next compared the density of interneurons within superficial and deep laminae and tested the independent and interactive effects of layer, species and area within each CaBP expressing interneuron subpopulation (Fig. 2d-f). Three-way ANOVA (species*area*layer) revealed significant main effects of layer (p = 9.4 x 10^-7^) and species (p = 4.2 x 10^-6^) on CR+ density, of layer (p = 2.6 x 10^-5^) and area (p = 0.0001) on CB+ density and of area (p = 0.0003) on PV+ density, but no interactive effects. Between-layer comparisons show a significantly greater density of CR+ neurons in the upper compared to deep layers for both species and areas (p < 0.0001 for all LSD post-hoc comparisons). Similarly, CB+ neuron is significantly greater in the upper compared to deep layers in mouse FC, mouse V1 and monkey V1 (p < 0.0001 for all LSD post-hoc comparisons). In contrast to CR+ and CB+ neurons, no significant between-layer differences were found for PV+ neurons.

Between-area comparisons revealed was no significant differences in the laminar density of CR+ neurons in either species (Fig. 2d). Interestingly, the laminar density of CB+ differed between areas in both species, within the upper layers (msV1 vs msFC, p= 0.00006; mkV1 vs mkLPFC, p = 0.0019; Fig. 2e). In addition, in mouse cortices, CB+ density in the deep layers was significantly greater in V1 than in FC (p = 0.047). PV+ neuronal density in the upper (msV1 vs msFC, p= 0.009; mkV1 vs mkLPFC, p = 0.023) and deep (msV1 vs msFC, p= 0.013; mkV1 vs mkLPFC, p = 0.033) layers was significantly greater in visual compared to frontal cortices of mouse and monkey (Fig. 2f).

Between-species comparisons revealed that compared to mouse, monkey V1 and LPFC both had significantly greater CR+ neuronal density in both the upper layers 1-3 (LSD post-hoc, msV1 vs mkV1: CR p = 0.014; msFC vs mkLPFC: CR p = 0.00045) and deep layers 4-6 (msV1 vs mkV1: CR p = 0.032; msFC vs mkLPFC: CR p = 0.0007; Fig. 2d). There were no between-species differences in the laminar-specific density of PV+ (Fig. 2f) and CB+ neurons (Fig. 2e) in either brain area.

The total and laminar density values observed in the present study for CR+, CB+ and PV+ interneurons in LPFC of n= 3 monkeys were highly consistent with the numbers observed in our previous study of the distribution of these neurons in the LPFC of a different cohort of six monkeys (Tsolias & Medalla, 2022) (Supplementary Material Fig. 2).

### Co-expression of CR+, CB+ and/or PV+ in individual neurons

We assessed the proportions, densities and laminar distributions of interneurons dual labeled for CRCB, CRPV and CBPV (Figs. 3 and 4). Two-way ANOVA (species*area) revealed significant between-species differences in the extent of CR+, CB+ and PV+ colocalization (total % colocalized main effect, species, p = 3.9 x 10^-6^), with no significant between-area differences. Specifically, the proportion of neurons exhibiting colocalization of these calcium binding proteins was significantly higher in mouse compared to monkey cortices (LSD post-hoc, msFC vs mkLPFC p=0.00001; msV1 vs mkV1 p=0.0002; Figs. 1b, 3a-c). This species difference was due to a greater proportion (Fig. 3c) and density (Fig. 4a, c) of CRCB and CBPV immunolabeled neurons in mouse compared to monkey (p<0.0001 for all comparisons). In the mouse, 3% and 3.7% of interneurons exhibited CRCB colocalization in V1 and FC respectively (Fig. 3c), and 4.4% and 5.7% of interneurons exhibited CBPV colocalization in V1 and FC respectively (Fig. 3c). In the monkey 0.18% and 0.56% of interneurons exhibited CRCB colocalization in V1 and LPFC respectively (Fig. 3c), and 2.32% and 1.04% of interneurons exhibited CBPV colocalization in V1 and LPFC respectively (Fig. 3c). Within each species and area, CBPV neurons were the most abundant type of neuron that co-expressed calcium binding proteins (Fig. 3c, 4c) being present at both a significantly higher density and proportion compared to CRPV and CRCB neurons (p < 0.01). Interestingly CRPV neurons were seen only in the monkey (Fig. 3c, 4b). Triple-expressing CR+CB+PV+ neurons were very rare (1-2 per case and only in monkey).

**Figure 3.**
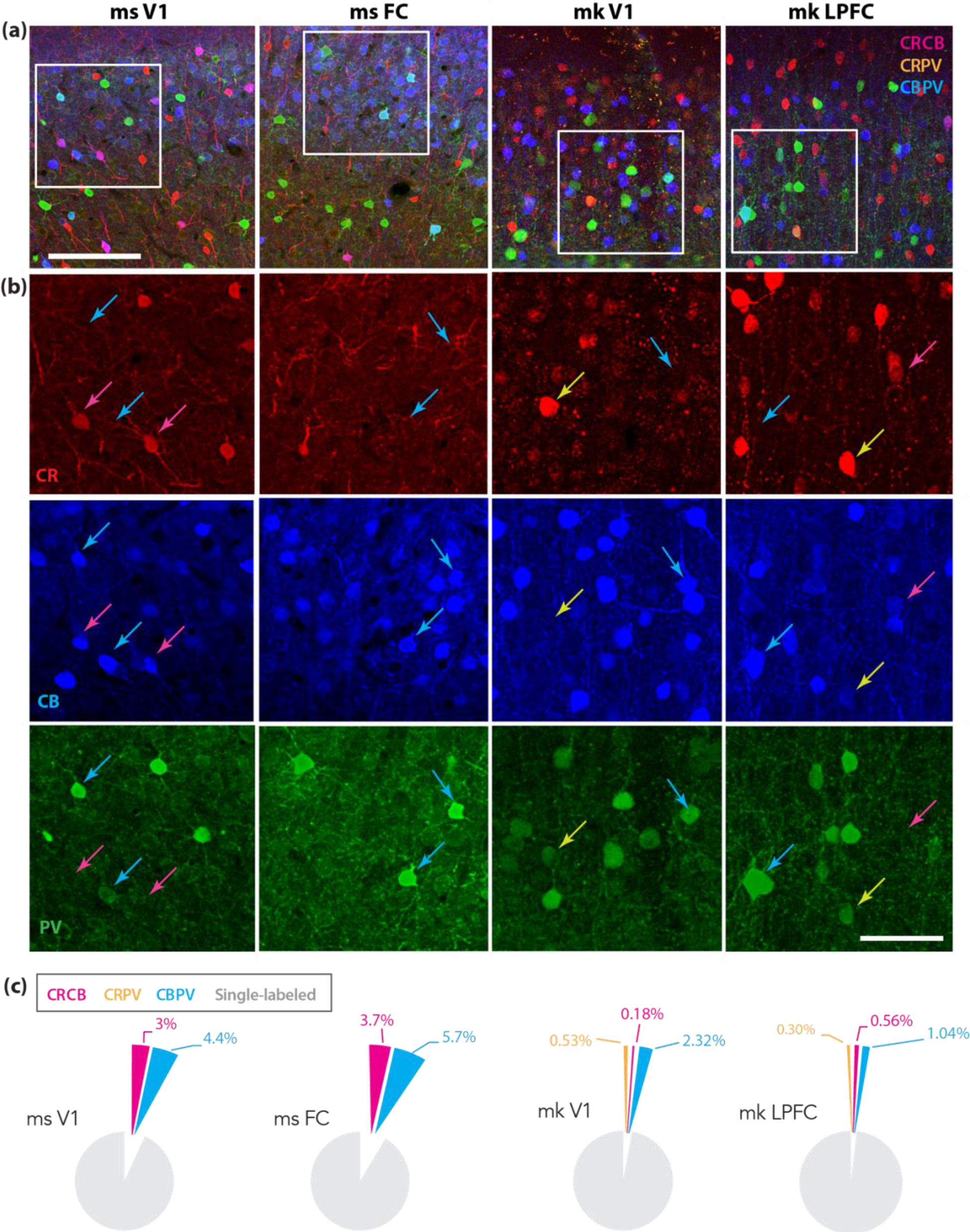
Co-localization of calcium binding proteins. a) Z-maximum projection confocal image stacks at low magnification (top, Scale bar: 100µm) and, **b)** High magnification images (boxed area in A; Scale bar: 50µm) with three channels separated, showing examples of neurons dual-labeled for CRCB (magenta arrows), CRPV (orange arrows) and CBPV (blue arrows). **c)** Pie charts showing percent of all interneurons co-expressing CRCB, CBPV and CRPV.

**Figure 4.**
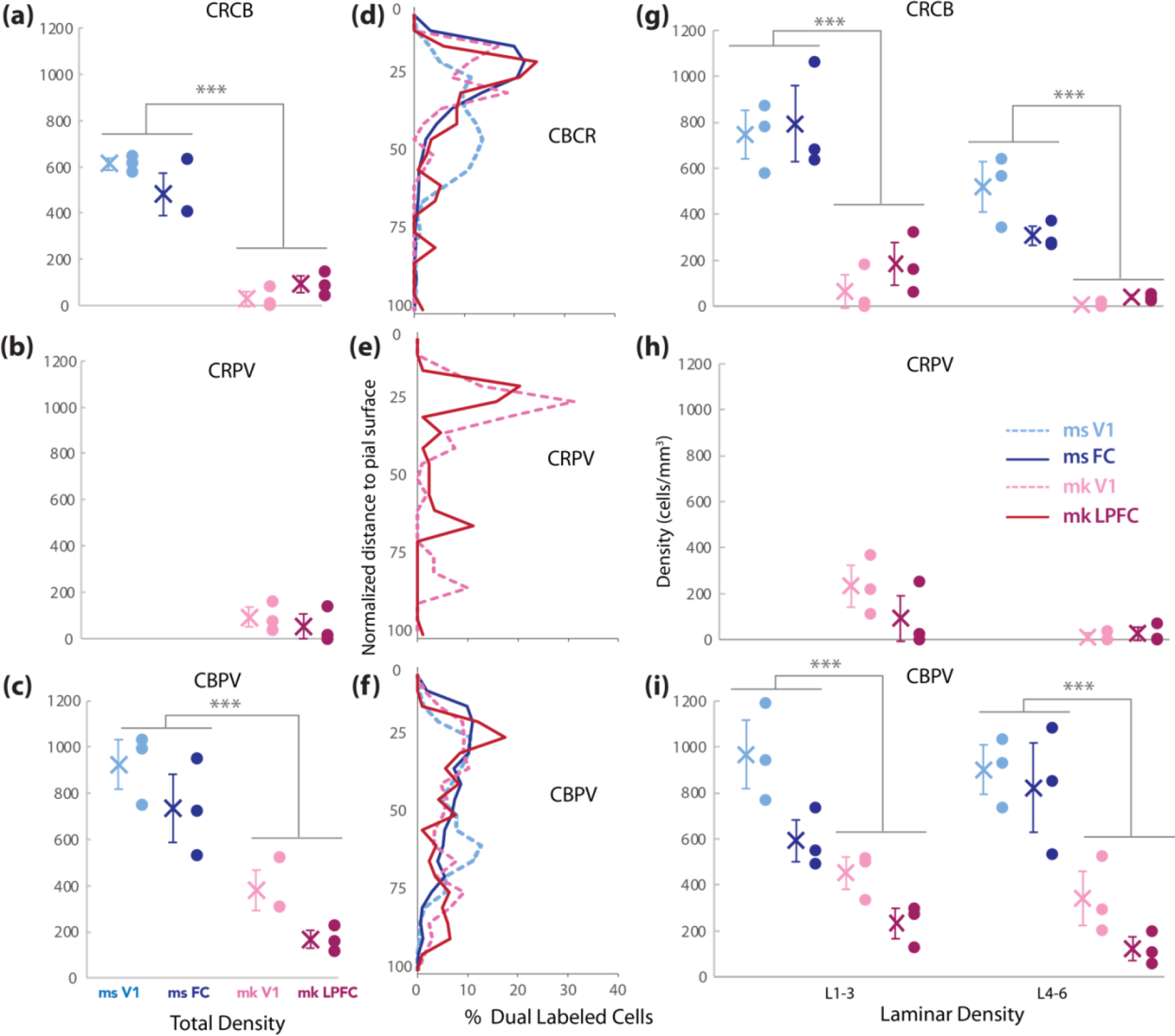
**Density and depth-distribution of calcium-binding protein co-expressing interneurons**. **a-c)** Total density, **d-f)** Cortical depth distribution and, **g-i)** Density in superficial (1-3) and deep (4-6) layers of interneurons co-expressing CRCB, CRPV or CBPV.

### Cortical depth and laminar distribution of neurons co-expressing CR+, CB+ and/or PV+

We assessed the distribution of CRCB, CRPV (present only in monkey), and CBPV neurons as a function of normalized cortical depth (Fig. 4d-f) and by quantifying their density in superficial layers and deep layers (Fig. 4g-i). The cortical depth distribution profiles of these calcium-binding protein co-expressing neurons were dependent on species and area (Fig. 4d-f). CRCB neurons were concentrated within the first 30% distance from pia in mouse FC and monkey V1 and LPFC (Fig. 4d). Mouse FC had significantly higher density of CRCB neurons within 10-20% cortical depth (approximately within layers 1-2) compared to mouse V1 and monkey LPFC (p<0.0001 for all comparisons; Fig. 4d). Furthermore, in mouse V1, a broader, deeper peak of CRCB neurons were observed, with a significantly greater density between 40-45% of cortical depth (p<0.05; Fig. 4d). CRPV interneurons, which were only present in monkeys, were concentrated within the most superficial 25-30% of cortical depth in both monkey areas (Fig. 4e). Both monkey areas showed a second, deeper peak in CRPV neurons, which was more superficial in LPFC (around 65% of cortical depth) compared to monkey V1 (around 85% of cortical depth; Fig. 4e). CBPV neurons were more evenly distributed across the cortical depth than were CRCB and CRPV neurons, (Fig. 4d-f). In mouse FC, CBPV neurons were present at significantly higher frequency within 5-15% of cortical depth (layer 1; p<0.05), but significantly lower frequency within 80-90% of cortical depth compared to monkey LPFC (p<0.05).

Consistent with the total regional densities, the densities of CRCB and CBPV neurons in both the upper and deep layers were significantly greater in mouse compared to monkey areas (Three-way ANOVA main effect species, CRCB p = 1.7 x 10^-8^; CBPV p = 3.8 x 10^-7^; p < 0.0001 for all LSD post-hoc comparisons; Fig. 4g-i). In addition, a significant main effect of layer (p = 0.0004) and a species*layer interaction (p = 0.025) were found for CRCB neuron density, which was higher in the upper compared to the deep layers of mouse, but not monkey cortex (Fig. 4g). CRPV neurons, which were rare and are only present in monkey, were also denser in upper compared to deep layers (main effect, layer p = 0.02; Fig 4h). Furthermore, a significant between-area laminar difference was found for CBPV neurons (main effect area, p = 0.004). Specifically, mouse V1 exhibited a significantly greater density of CBPV neurons in the upper layers compared to mouse FC (LSD post-hoc, p = 0.02; Fig. 4i).

### Somato-dendritic properties of CR+, CB+, PV+ neurons

Inhibition within cortical circuits is determined not only by the distribution of different classes of interneurons but also by their morphological features. Thus, we reconstructed the soma and dendrites of CR+, CB+, PV+ neurons (*n*=386) with somata in L2-3 (Fig. 5).

**Figure 5.**
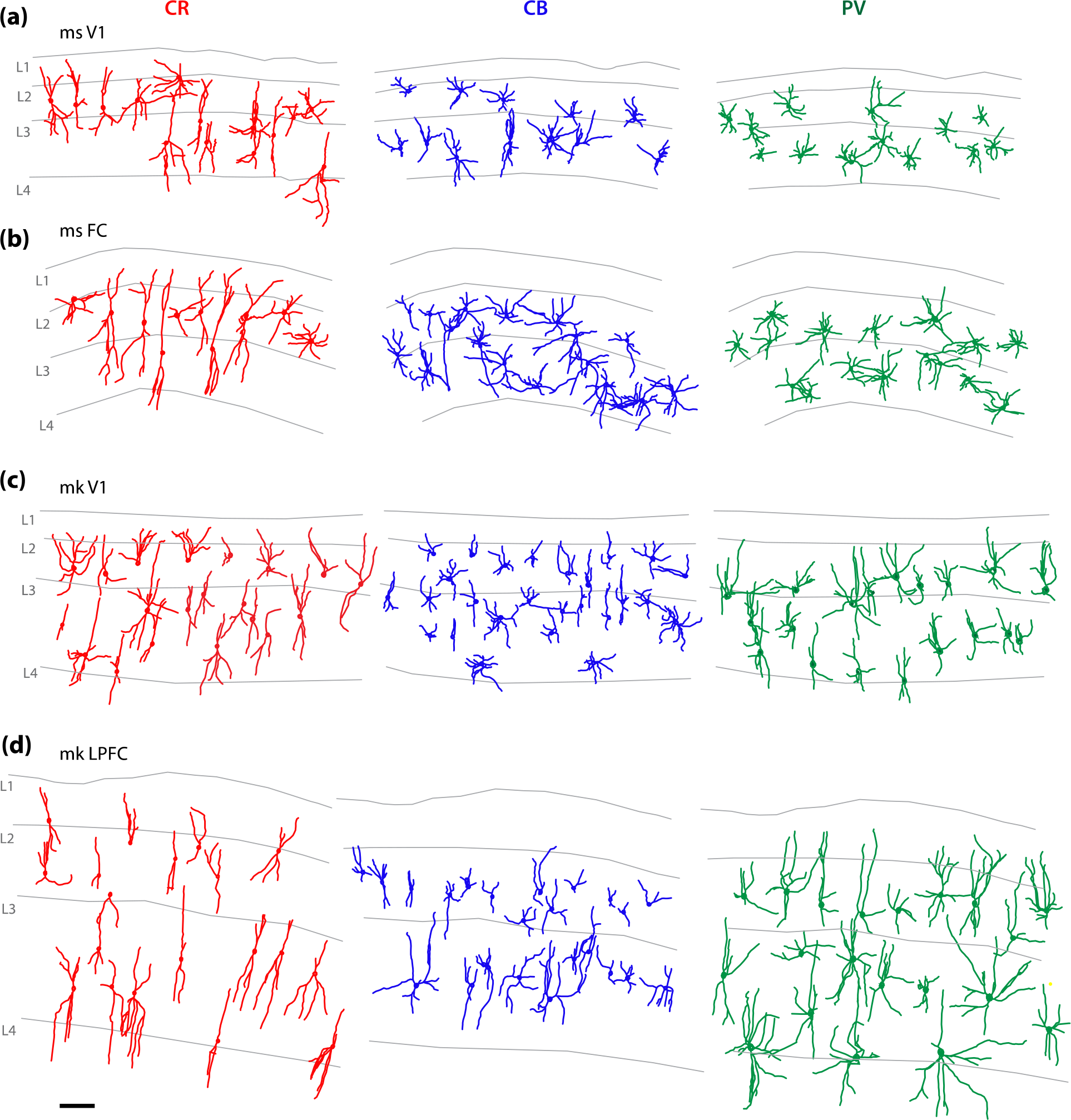
3D tracings of interneuron somata and dendrites for morphological analyses. Representative examples of reconstructed CR+ CB+ and PV+ interneurons (superimposed data from multiple sections) in **a)** mouse V1, **b)** mouse FC, **c)** monkey V1 and, **d)** monkey LPFC. Labels indicating layers 1-4 (L1-4) are placed on the top boundary of each layer. Scale bar: 100 µm.

Three-way ANOVA revealed significant main effects (Species, *p* = 0.00000007; CaBP *p* = 5.7 x 10^-18^) and interactive effects (species*CaBP, *p* = 0.03) of species and CaBP-expressing cell type on soma volume (Fig 6a; see Supplementary Material Table S1). For CR+ and PV+, but not CB+, interneuron subtypes, soma volume was significantly greater in monkey compared to mouse (Fig. 6a). Significant differences in soma volume between the different CaBP expressing interneuron subtypes were also found consistently within species. Within both species, CB+ neurons in both brain areas had significantly smaller soma volume compared to PV+ neurons (*p* < 0.001 for all comparisons). PV+ neurons had the largest soma volumes, significantly larger than both CR+ and CB+ neurons (*p* < 0.01 for all comparisons), consistent with work from previous studies (Gabbott & Bacon, 1996a; Glezer et al., 1998; Sherwood et al., 2007). CR+ neurons were intermediate in size, having smaller volumes than PV+ neurons in both species and areas (*p* < 0.01 for all comparisons), and larger volumes than CB+ in monkey V1 (CB vs CR *p* = 0.001) and LPFC (CB vs CR *p* = 0.008).

**Figure 6.**
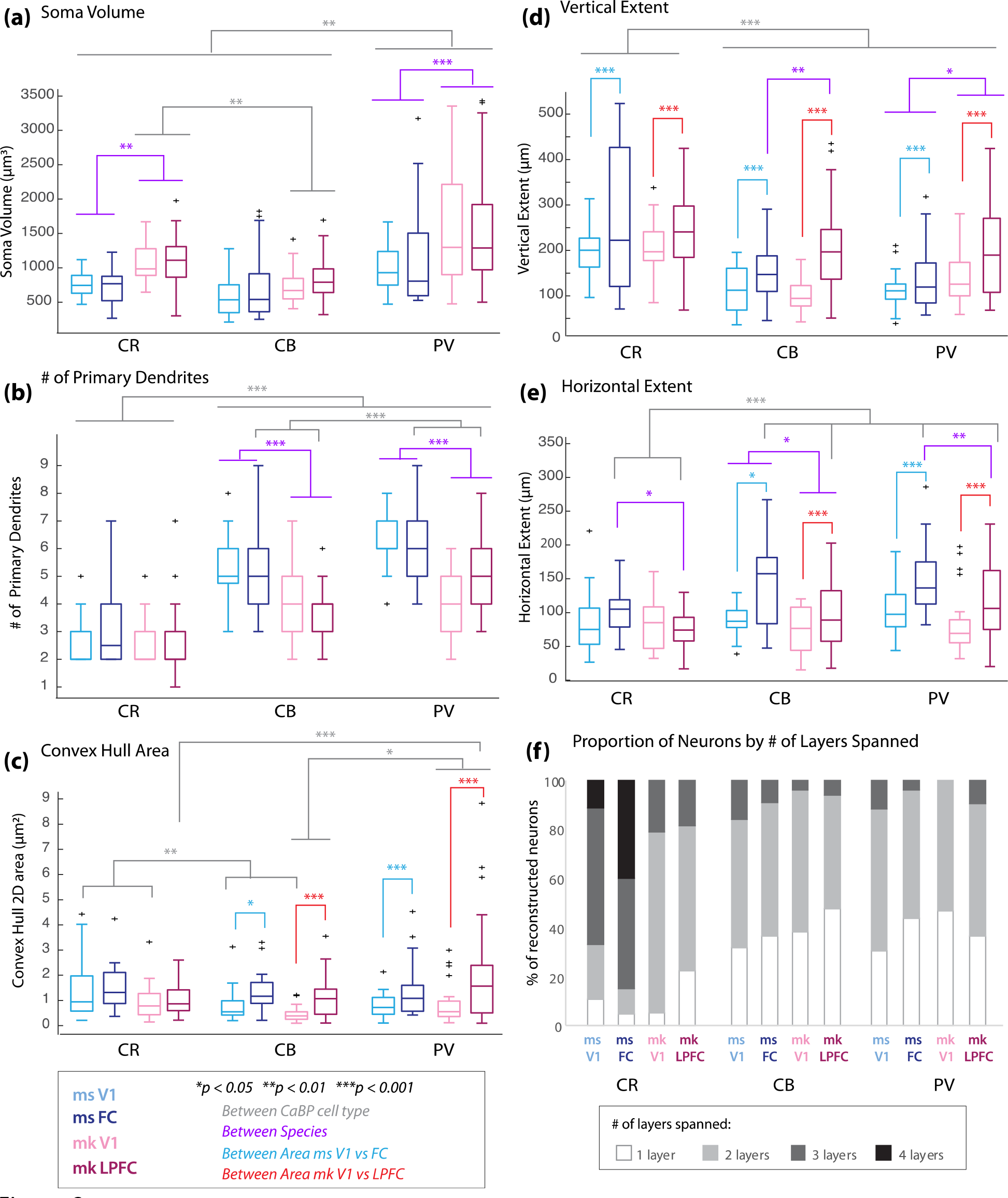
Quantification of somato-dendritic morphological features of CR+ CB+ and PV+ interneurons. Box and whisker plots showing: a) Soma volume, b) Number of primary dendrites; **c)** 2D convex hull area of coverage of dendritic arbors, **d)** Maximum vertical extent and, **e)** Maximum horizontal extent of dendritic arbors. Brackets delineate significant differences between groups based on three-way ANOVA Post-hoc, Fisher’s LSD analyses. **f)** Proportion of neurons from each subtype by the number of layers spanned by dendritic arbors.

A significant main effect of species (*p* = 1.15 x 10^-15^) and CaBP cell type (*p* = 4.76 x 10^-48^) was also found for the number of primary dendrites and a significant interaction between species and CaBP (*p* = 0.0002; Fig. 6b). Within each species and area, both CB+ and PV+ neurons had a higher number of primary dendrites compared to CR+ neurons (*p* < 0.001 for all comparisons, Supplementary Material Table S1; Fig. 6b). However, between-species differences were found only for CB+ and PV+ interneurons: for CB+ and PV+, but not CR+ neurons, there was a higher number of primary dendrites in mouse V1 and FC compared to monkey V1 and LPFC (*p* < 0.001 for all comparisons, Supplementary Material Table S1; Fig. 6b).

We then further characterized dendritic morphological features, quantifying the total 2D convex hull dendritic area coverage, vertical and horizontal spans, and polarity of interneurons across species, cortical area and CaBP expressing cell types. The total 2D convex hull area of interneurons exhibited significant differences across cortical areas (*p* = 8.04 x 10^-7^) and CaBP expressing cell type (area*CaBP interaction, *p* = 0.007), but no significant main effect of species (Fig. 6c; Supplementary Material Table S1). Specifically, CB+ and PV+ neurons, but CR+ neurons exhibited smaller convex hull area span in V1 than in frontal areas in both mouse and monkey (Fig. 6c; p < 0.02 for all comparisons). Between-CaBP cell type differences depended on area, with CB+ neurons being smaller than PV+ neurons in monkey frontal and visual cortices (p < 0.01 for all comparisons), but smaller than CR+ neurons only in visual, but not frontal cortex in both monkey and mouse (p < 0.05 for all comparisons; Fig. 6c).

We next assessed the vertical (Fig. 6d) and horizontal (Fig. 6e) dendritic extent of each cell and found a significant main effect of species (horizontal, *p* = 0.000016; vertical *p* = 0.013), area (horizontal *p* = 3.63 x 10^-9^; vertical *p* = 1.32 x 10^-11^) and CaBP-expressing cell type (horizontal *p* = 0.000017; vertical *p* = 1.83 x 10^-15^). There was a significantly greater vertical dendritic extent of interneurons in the frontal compared to visual cortices in both species (main effect area, *p* = 1.32 x 10^-11^; Fig. 6d), while a significantly greater horizontal extent was found for mouse compared to monkey, in both cortical areas (main effect species, *p* = 0.000016; Fig. 6e).

CB+ and PV+ neurons had consistently larger horizontal extents, but smaller vertical extents compared to CR+ neurons in both areas and species (main effect CaBP cell type, horizontal *p* = 0.000017; vertical *p* = 1.83 x 10^-15^; Fig. 6d, e). For vertical extents, this between-cell type difference was additionally dependent on species, whereby CB+ and PV+, but not CR+ neurons exhibited smaller vertical dendritic spans in mouse than in monkey (Species*CaBP cell type interaction, *p* = 0.02, all LSD post-hoc *p* < 0.01, Supplementary Material Table S1; Fig. 6d). In contrast, between-CaBP expressing cell type difference in horizontal span was additionally dependent on area, with CB+ and PV+ but not CR+ neurons exhibiting greater horizontal extents in frontal than in visual cortices in both species (area*CaBP cell type interaction, *p* = 0.002, all LSD post-hoc *p* < 0.01, Supplementary Material Table S1; Fig. 6d).

We quantified the number of layers spanned by the dendritic arbor of each cell and assessed the proportion of neurons with a given laminar span (Fig. 6f). We found that compared to CB+ and PV+ neurons, there was a greater proportion of CR+ neurons with dendrites that spanned 3 or more layers and a lower proportion of CR+ neurons with dendrites contained within 1 layer (ms V1 *X^2^* = 24.1, p = 0.0005; ms FC *X^2^* = 49.1, p = 7.09 x 10^-9^ ; mk V1 *X^2^* = 15.8; p = 0.003; mk LPFC *X^2^* = 9.3; p = 0.052; Fig. 6f). Further, within the population of CR+ neurons, there was a species difference in distribution of cells by laminar span. Specifically, in mouse V1 and FC, the majority CR+ neurons had dendrites spanning 3 or 4 layers, in higher proportions than monkey CR+ neurons (*X^2^* = 54.4, p = 1.5x x 10^-8^; Fig. 6f). This species difference in laminar span was not present in CB+ and PV+ subpopulations; these neurons had dendrites constrained to a single layer (the same layer as their cell bodies) or spanning only 2 layers (crossing one laminar boundary) in both mouse and monkey areas.

We then assessed the morphology of dendrites based on polarity, namely as multipolar or bipolar type 1 or bipolar type 2 subtypes (Fig. 7a). Multipolar neurons are characterized by multiple primary dendritic branches emerging across the extent of the soma surface. Bipolar neurons by contrast exhibit either two (bipolar type 1) or, more rarely, multiple (bipolar type 2) primary dendrites emerging from opposite poles of the soma. A quantitative measure of polarity was obtained by plotting a polar angular histogram of dendritic length, then calculating the vector [x, y] projection of the polar plot (Fig. 7b, c), as described previously (Guillamon-Vivancos et al., 2019). Analyses of polarity revealed significant main effects of species (*p* = 1.18 x 10^-8^) and CaBP cell type (*p* = 0.019) on the Y (vertical) component of the polar plot vector projection (Fig. 7b, c). Specifically, CB+ and PV+ neurons exhibited between-species differences in polarity, with monkey neurons showing greater vertical polarization than mouse neurons in both brain areas (*p* < 0.0001), while CR+ showed greater vertical polarization only in monkey LPFC compared to mouse FC (*p* = 0.007; Fig. 7c; Supplementary Material Table S1).

**Figure 7.**
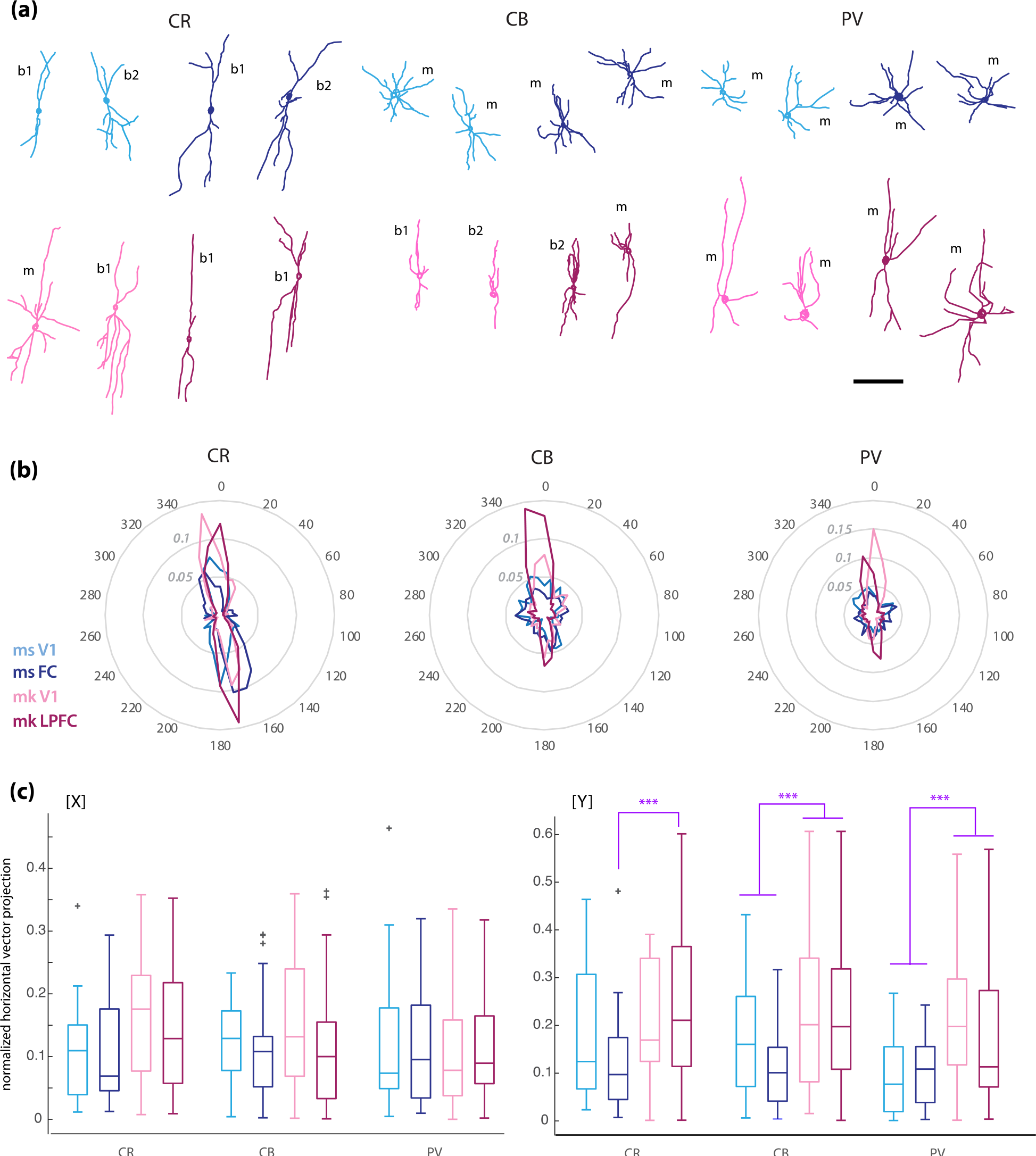
Quantification of polarity and orientation of CR+, CB+ and PV+ interneuron dendrites. **a)** Representative reconstructions of cells exhibiting bipolar type 1 (b1) or type 2 (b2), or multipolar (m, CB and PV only) morphologies from each subclass. **b)** Radial histogram of normalized dendritic length (circles and light gray font) within angular bins (dark gray font), averaged for CR+ CB+ and PV+ neurons for each area and species. **c)** Box and Whisker plots of the “X” and “Y” projections of the summation vectors of the polar histograms.

### Dimensional reduction analyses reveal clustering of CR+ CB+ PV+ expressing neurons based on multivariate somato-dendritic morphological features

Our data revealed significant differences in specific morphologic features of superficial layer interneuron subclasses that seem to be mainly dependent on CaBP expression, with significant interactions with species and cortical area. We further adressed whether the combination of distinct somato-dendritic morphological features can define populations of inhibitory neurons using cluster analyses and multidimensional scaling methods (Medalla et al., 2020). Thus, we assessed the relative similarity and clustering of neurons in each group by conducting non-metric multidimensional scaling (NMDS) based on pairwise distance correlations across the multivariate set of 7 somato-dendritic outcome measures (see Methods) (Fig. 8). This analysis yielded clusters of individual neuron data points based on a proximity distance matrix, such that points that are closer together are more similar with respect to the set of morphological features (see also Dombrowski et al., 2001; Gilman et al., 2017; Hsu et al., 2017). To further validate the unsupervised clustering and NMDS results, MANOVA was performed and the outcome measures that influenced classification based on species, area and CaBP cell type were determined. The NMDS plot based on somato-dendritic morphology revealed a visible clustering of neurons from the 12 subgroups based mainly on CaBP cell type, and to a much lesser extent, species (Fig. 8a-c, left). Annotation of cells by CaBP cell type revealed that PV+ and CR+ neurons are morphologically distinct from each other, forming clear segregated clusters (Fig. 8c). In contrast, the CB+ neuron cluster overlapped with both the CR+ and PV+ neuron clusters. MANOVA analyses validated these findings, showing how the 12 subgroups were significantly clustered into 6 groups based on six significant canonical variables that delineated between species and CaBP cell type (Fig. 8a- c, right; Supplementary Material Fig. S3, Table S2). The first canonical variable separated two main groups based on CaBP, with the first main cluster consisting of CR+ neurons (and some CB+ cells), and the second main cluster consisting of both CB+ and PV+ neurons (Fig. 8a, right; Supplementary Material Fig. S3). The rest of the canonical variables separated between species (Supplementary Material Fig. S3). The absolute value of the coefficients of the canonical variables determines the strength of each outcome measure as a predictor of group separation. Among the outcome measures, the number of primary dendrites and vertical extents had the strongest influence (the largest 1^st^ canonical coefficients) on the first discriminator, which separated CR+ neurons from CB+/PV+ neurons (Supplementary Material Fig. S3, Table S2). Subsequently, soma size, the number of layers spanned and horizontal extents were the strongest between-species discriminators (2^nd^-6^th^ canonical coefficients; Supplementary Material Table S2).

**Figure 8.**
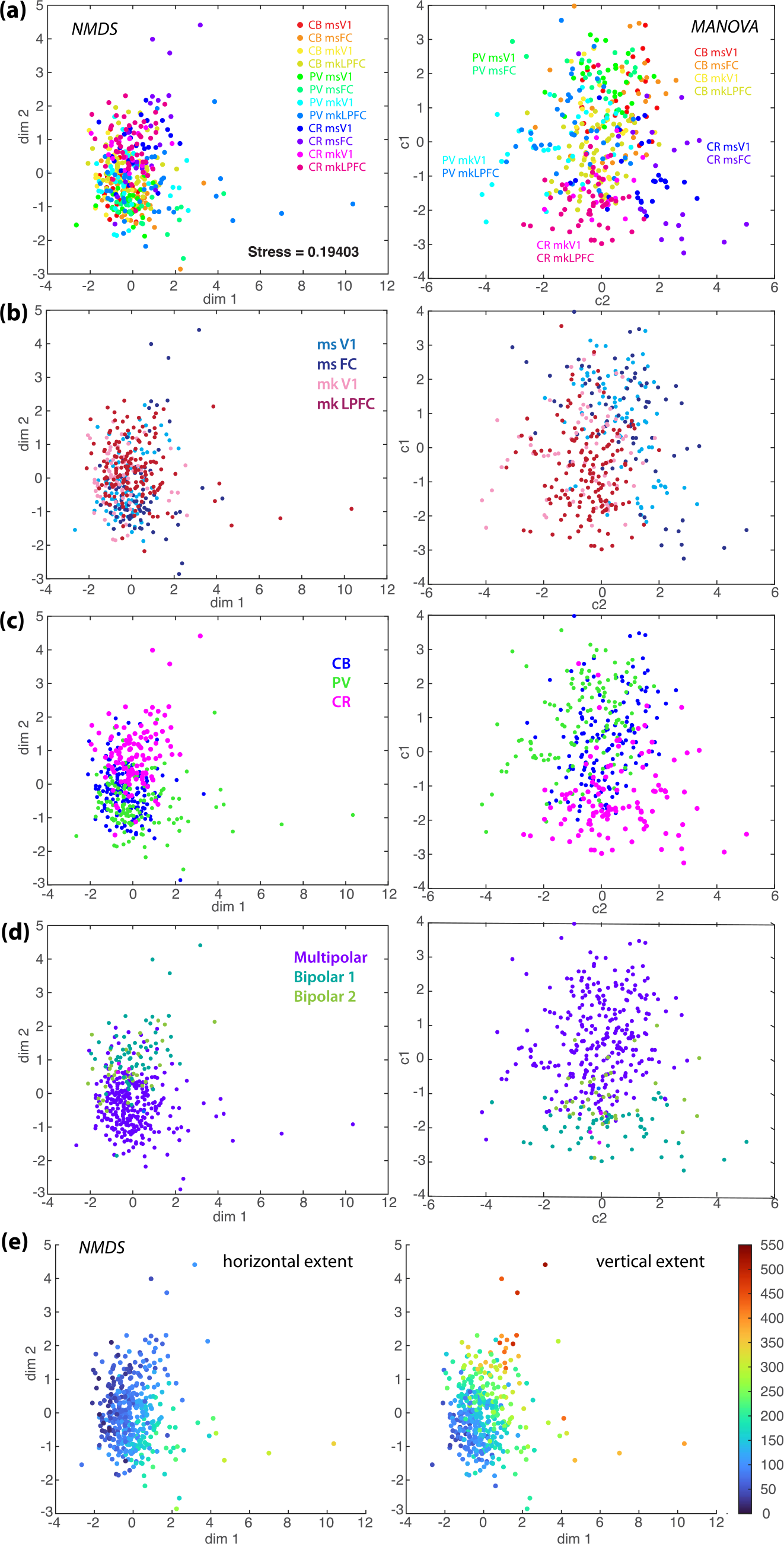
Dimensional reduction and clustering of total interneuron population based on multivariate morphological features. a-d) NMDS plots (left) showing clustering of individual neurons based on dendritic morphological features, validated with supervised cluster plots based on MANOVA (right). Individual neurons (data points) on the plot are annotated by: **a)** Experimental group by species x area x CaBP; **b)** Species x area showing clustering based on species but not area; **c)** CaBP expression, showing a strong segregation of CR+ and PV+ neurons with CB+ neurons being intermingled between the two other populations of interneurons; **d)** Morphological classification showing segregation of bipolar (type 1 and type 2) and multipolar neurons. **e)** The same NMDS clustering plot as *a-d*, but data points are annotated based on a gradient map of horizontal (left) and vertical (right) extents. Note that neurons with large vertical but small horizontal extents (top part of the plot) correspond to CR+ neurons in *c* and bipolar neurons in *d*.

We further annotated the clusters based on morphological classes and found that the NMDS and MANOVA clustering corresponded with a separation across multipolar versus bipolar subclasses of interneurons (Fig. 8d). When looking at the NMDS plot with a gradient map of the vertical and horizontal extent of each neuron, we find that neurons with larger vertical extents, but smaller horizontal extents clustered together (Fig. 8e). Consistent with our ANOVA comparisons, the sub clustering based on morphological features seems to be independent of laminar location (of somata in L2 versus L3 or on distance from pia; data not shown).

Clustering of interneurons based on multivariate analyses showed that CaBP is a strong determinant of dendritic morphological subclasses. Thus, we further examined if neuronal subclasses were present within populations of CR+ CB+ and PV+ by running NMDS analyses on each of these populations separately (Fig. 9). Within each CaBP cell type, the degree of separation based on species and morphology differ. CB+ and PV+ but not CR+ neurons exhibited subclusters based on species (Fig. 9a). However, there was no separation between areas, indicating that the somato-dendritic morphology of interneurons from visual and frontal cortices is relatively similar within each species and CaBP cell type. Interestingly the NMDS clustering plots of CR+, CB+ and PV+ neurons corresponded to distinct bipolar versus multipolar morphologies (Fig. 9B). CR+ interneurons were defined by segregated bipolar (type 1 and 2) versus multipolar neuron subclusters. These subclusters of morphologically distinct CR+ neurons correspond to two groups with distinct vertical and horizontal extents; CR+ bipolar cells have large vertical but small horizontal extents, while CR+ multipolar neurons have small vertical and horizontal extents (Fig. 9C, D).

**Figure 9.**
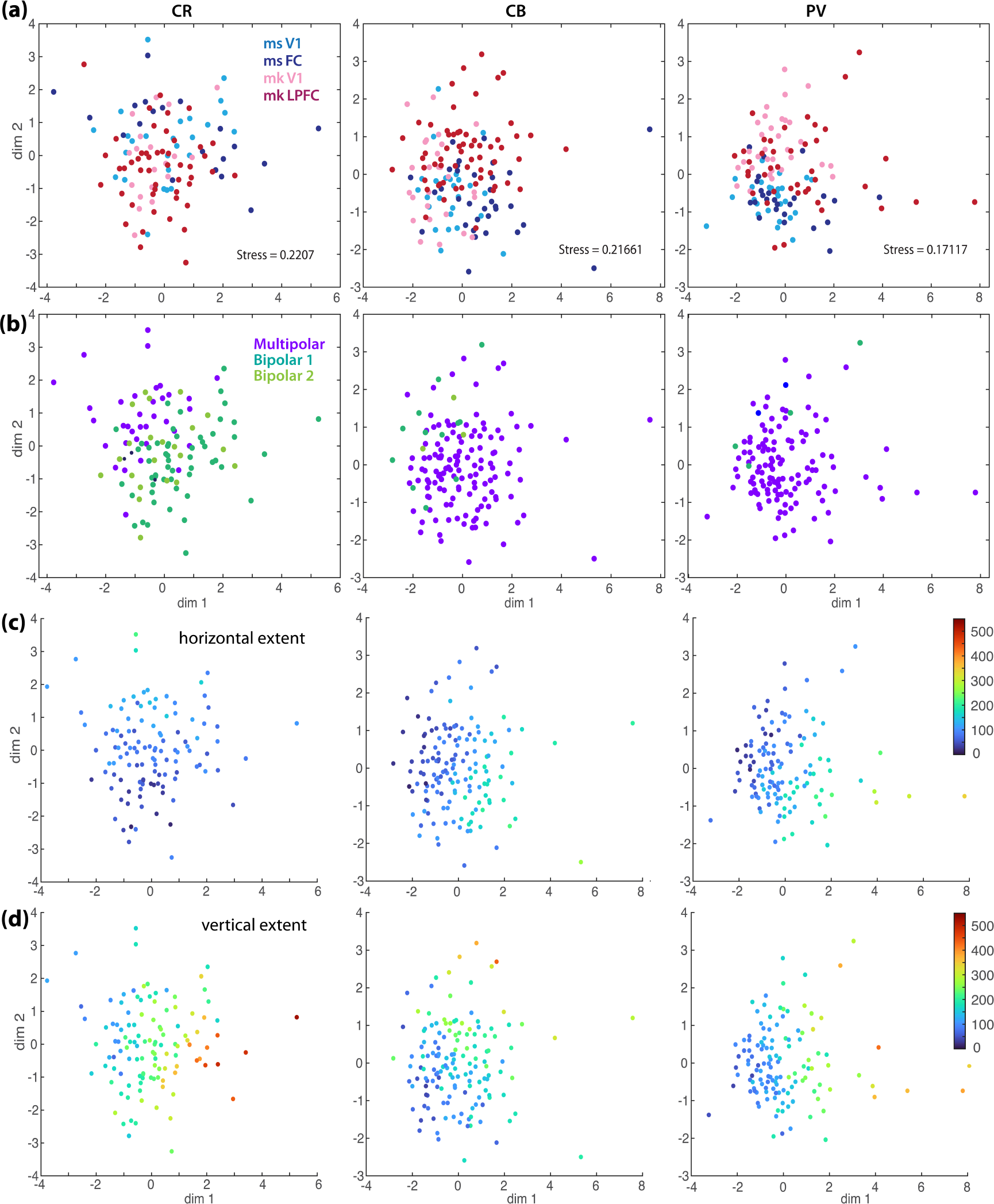
Dimensional reduction and clustering of neurons within the CR+, CB+, PV+ subpopulations based on multivariate morphological features. a) Separate NMDS plots of clustering of CR+, CB+ and PV+ individual neurons based on dendritic morphological features, with individual neurons (data points) on the plot annotated by species x area. Note that clustering based on species, but not area is dependent on CaBP subtype. CR+ neurons showed no clustering based on species, while CB+ and PV+ neurons showed separation by species; **b)** The same NMDS plots as *a*, but with data points annotated by morphological classification showing a segregation of bipolar and multipolar neurons, which depended on CaBP subtype. CR+ neurons exhibited more bipolar morphology, while PV+ neurons were almost exclusively multipolar. CB+ neurons are primarily multipolar with a smaller subset having bipolar morphology. **c, d)** The same NMDS clustering plots as *a-b*, but data points are annotated based on a gradient map of horizontal (c) and vertical (d) extents. Note that CR+ neurons have large vertical but small horizontal extents, while PV+ neurons have large vertical and horizontal extents consistent with their wide arbor multipolar morphologies.

In contrast to CR+ interneurons, the delineation between bipolar versus multipolar neurons was less pronounced in CB+ neurons and was not present in PV+ neurons (Fig. 9B). Most CB+ and almost all PV+ neurons were multipolar. However, there was a small subcluster of CB+ interneurons that were bipolar in morphology (Fig. 9B). Interestingly, visualization of the horizontal and vertical extent of individual CB+ and PV+ neurons revealed the presence of small arbor (small vertical and horizontal extents) versus wide arbor (large vertical and horizontal extents) multipolar neurons respectively (Fig. 9C, D).

In summary, these data show that CR+ neurons have dendritic morphologies that are markedly distinct from PV+ and CB+ neurons in both species and areas. In contrast, PV+ and CB+ interneurons in both species and areas were morphologically more similar to each other. Furthermore, CR+ neurons represented at least two morphological subclasses -multipolar and bipolar- across species and areas. However, CR+ neurons did not generally exhibit significant between-species and between-area differences in dendritic morphology. In contrast CB+ and PV+ inhibitory neurons exhibit a modest but statistically significant degree of between-species differences with regards to somato-dendritic morphology, but no regional differences across visual and frontal cortical areas.

## DISCUSSION

### Summary of key findings

In ongoing efforts to understand how the diverse cellular constituents of local circuits may contribute to area- and species-specific cortical cytoarchitecture and functions, it is essential to quantify how the distribution and features of both excitatory pyramidal neurons and inhibitory interneurons vary across brain areas within and between species. At opposite ends of the rostro-caudal neuroaxis, and with distinctive cytoarchitecture and functions across phylogeny (review: Pandya, 1985; Yeterian et al., 2012), the primary sensory visual cortex (V1) and the frontal cortex in mouse (FC) and lateral prefrontal cortex in monkey (LPFC) are ideal brain regions in which to perform comparative studies such as these. We and others have established that individual pyramidal neurons scale dramatically in size from the primary visual cortex to the frontal cortical regions in the primate but not in the mouse (e.g. Ballesteros-Yanez et al., 2007; Elston et al., 2011; Elston & DeFelipe, 2002; Gilman et al., 2017) (reviews:Elston et al., 2011; Luebke, 2017; Wittenberg, 2012); whether inhibitory interneurons show area or species-specific features - particularly with regard to their structure- is considerably less well understood. Here, we directly compared the areal and laminar distribution and detailed somato-dendritic morphological features of 3 major classes of neocortical interneurons identified by the presence of the calcium binding proteins (CaBPs) -CR, CB and PV- in the mouse V1 and FC versus the monkey V1 and DLPFC. Consistent with previous studies, we found that these distinct CaBP expressing interneuron types are largely non-overlapping populations and are distinct with respect to their laminar distribution and somato-dendritic morphology (reviewed in DeFelipe, 1997; DeFelipe et al., 2013; Tremblay et al., 2016). Specifically, our data confirms that: 1) CR+ and CB+ interneurons are concentrated within superficial layers 1-3, while PV+ interneurons are more generally distributed uniformly across deeper layers 2-4 in both areas and in both species; 2) There is a significantly higher proportion of interneurons that co-express two different CaBPs in the mouse compared to monkey; 3) The density and proportion of CR+ neurons are higher in monkey than in mouse cortices, and; 4) There is a modest degree of scaling of interneurons with regard to soma volume, with CR+ and PV+ neurons being slightly larger in monkey than in mouse.

This study produced novel information related to the interactive effects of species and area on the distribution and somato-dendritic morphology of interneurons, which are dependent on their CaBP expression. Specifically, we demonstrate that: 1) There are areal differences in CB+ and PV+ neuron densities, with significantly higher densities in V1 compared to frontal areas in both species. By contrast the density of CR+ neurons do not differ between areas in either species; 2) The relative proportions of the 3 populations of interneurons differ between areas and species. While CB+ neurons comprise the majority of the interneuron population in mouse and monkey V1 and in mouse FC (44%, 50% and 42% respectively), in monkey LPFC the predominant class is CR+ (40%), with CB+ comprising the second most abundant type (35%); 3) The dendritic morphology of CR+ neurons are markedly distinct from PV+ neurons and to a lesser extent from CB+ neurons in both species and areas; 4) While the CR+ neuron proportion and densities are greater in monkeys compared to mouse for both areas (but most notably in LPFC), CR+ neurons do not exhibit significant between-species and between-area differences in dendritic morphology. In contrast we observed a modest degree of between-species differences with regard to somato-dendritic morphology of CB+ and PV+ inhibitory neurons.

### Comparative proportions of CR+ CB+ and PV+ interneurons

The key difference between species was the significantly higher proportion of CR+ interneurons in monkey compared to mouse in both visual and frontal cortices. This is consistent with previous studies showing an expansion in the proportion of CR+ interneurons in monkey neocortex, that is believed to account for the significantly higher number of GABAergic interneurons in the primate (Dzaja et al., 2014; Hladnik et al., 2014). This finding of higher numbers and density of CR+ interneurons in the monkey was more pronounced in frontal compared to visual areas; In monkey LPFC the most abundant type of interneuron was CR+, which was 2-3x greater in density compared to CR+ neurons in mouse FC. CR+ neurons have been shown to be an interneuron population uniquely expanded in neurogenesis of primate-specific association cortices (Fogarty et al., 2007; Gonchar et al., 2007; Ma et al., 2013; Radonjic et al., 2014). These neurons are postulated to play a role in the function specifically of association cortices that are much more developed in primates than in rodents (Dzaja et al., 2014; Ma et al., 2013). By virtue of their inhibitory connections to other GABAergic neurons in the supragranular layer in primates (Dzaja et al., 2014; Hladnik et al., 2014; Meskenaite, 1997), CR+ interneurons participate in ‘disinhibitory” functions, adding more control and complexity to inter-areal and intra-areal processing that may be necessary for the integrative functions of these higher association areas (Dzaja et al., 2014).

Our study confirmed that a small number of inhibitory interneurons express more than one CaBP in both species and in both V1 (Kooijmans et al., 2020) and frontal regions. As reported earlier (Kooijmans et al., 2020), the proportion of dual-labeled neurons is significantly higher in the mouse than in the monkey V1 (7.4% mouse, 2.93% monkey). Similarly, there is considerably more dual labeled CaBP+ neurons in mouse compared to monkey frontal areas (9.4% mouse, 1.9% monkey). This is interesting, given that primate interneuron development may represent more highly divergent neurogenetic pools compared to the rodent (Fogarty et al., 2007; Gonchar et al., 2007; Ma et al., 2013; Radonjic et al., 2014). Of note, in the mouse all dual labeled neurons were either CBCR+ or CBPV+, with the majority falling into the latter category. In monkey V1 and LPFC, but not in mouse, a small number of CRPV+ neurons were also observed. Whether these CRPV+ interneurons represent a unique class of rare primate-specific neurons is a question that requires further investigation.

A recent study comparing interneuron proportions using the same markers in mouse and monkey V1 (Kooijmans et al., 2020) reported different findings than those reported here. These investigators found only modest differences in the proportions of the 3 classes of interneurons in the two species- the predominant type was PV+ (42% mouse, 48% monkey) followed by CB+ (24% mouse, 29% monkey) then CR+ (24% mouse, 20% monkey). In our assessment of the relative proportions of CaBP+ interneurons we found that the CB+ subtype predominated in both mouse V1 and monkey V1, with the second most abundant type being PV+, followed by CR+. With regard to monkey LPFC, the proportions of CR+ CB+ and PV+ neurons in LPFC were consistent with numbers reported in (Conde et al., 1994), and the densities of CB+ and PV+ neurons was similar to that reported in (Dombrowski et al., 2001). However, (Dombrowski et al., 2001) report a lower density of CR+ interneurons than reported here. These discrepant results point to the well-known difficulty in comparing quantitative findings related to interneuron markers across studies. Methodological differences such as tissue preparation, antigen retrieval, specific immunohistochemical procedures and antibodies used, can impact the numbers of neurons labeled (see Winsky & Kuznicki, 1996). Nevertheless, when tissues are processed in a near identical manner across brain areas and species as in the present study and in others from our group (Tsolias & Medalla, 2022), comparisons of distributions and somato-dendritic morphologies are likely reliable.

### Laminar distribution of inhibitory neurons expressing CR, CB or PV

Analysis of the superficial (layers 1-3) versus deep (layers 4-6) laminar distribution of the 3 classes of interneuron revealed differential patterning between areas and species. The laminar density of CB+ differed between areas in both species, especially within the upper layers with mouse V1 showing the highest CB+ density in upper layers. In monkey cortices, CB+ density in the deep layers was significantly greater in V1 than in LPFC. Similarly, PV+ neuronal density in the upper layers was significantly greater in visual compared to frontal cortices of both mouse and monkey. Further, mouse V1 showed significantly higher PV+ density in deep layers compared to mouse FC, while no significant between-area difference in PV+ density in the deep layers was found in the monkey. While there was no difference in the laminar density of CR+ neurons between areas, significant between-species differences were found. As expected, based on the higher total density of CR+ neurons, and their preferential distribution in superficial layers, there was a higher density of CR+ neurons in the upper layers of both visual and frontal areas in monkey compared to mouse. Between species differences in CB+ and PV+ laminar density were less pronounced, with only PV+ neuron density in the upper layers showing a significantly higher density in monkey V1 compared to mouse V1. Taken together with their different proportions and known differences in postsynaptic targets, these differences in laminar location of diverse populations of highly target-specific interneurons likely confer distinct excitatory:inhibitory synaptic processes in different brain areas and regions. Additional studies including computational network modeling and optogenetic cell-specific circuit manipulations are required to understand these nuanced regional and species specializations in distribution of diverse interneurons.

### Single-cell somato-dendritic morphological features of interneuron subtypes

Dimensional reduction and clustering of the total interneuron population based on multivariate morphological features showed that the somato-dendritic morphology of L2-3 inhibitory interneurons is more dependent on CaBP expression than on species and area. This confirms previous work showing that CR+ CB+ and PV+ neurons are morphologically distinct subtypes (Buzsaki et al., 2004; Conde et al., 1994; DeFelipe et al., 2013; Monyer & Markram, 2004; Yuste, 2005; Zaitsev et al., 2005; Zaitsev et al., 2009). Our quantitative analyses revealed how these three cell types cluster based on their morphological features. Specifically, CR+ and PV+ neurons appear to be highly morphologically distinct populations, forming distinct clusters of neurons. The CR+ population is largely comprised of bipolar neurons with long vertical extents and strong vertical polarization. In contrast, PV+ neurons are almost exclusively multipolar neurons with large horizontal and moderate vertical extents. Interestingly, CB+ neurons are somewhat intermingled between clusters of CR+ and PV+ neurons- the largest cluster was comprised of those with multipolar morphology and a smaller subcluster was comprised of bipolar neurons. This is consistent with our finding that both CB+ and PV+ neurons had a higher number of primary dendrites and larger horizontal extents but smaller vertical extents compared to CR+ neurons. Accordingly, a larger proportion of CR+ neurons span more than 1 layer compared to the other CaBP-expressing subtypes, especially in mouse areas. PV+ neurons, which also had the largest soma volumes of all subtypes, had the largest within-layer horizontal spans, consistent with the large basket cell morphology associated with PV+ neurons in other studies (Jones, 1986; Markram et al., 2004). These findings suggest that individual L2-3 PV+ and CR+ neuron dendrites receive distinct laminar inputs, with CR+ neurons integrating inputs from different layers and PV+ neurons integrating inputs within a given lamina.

Interestingly, based on dendritic topology and soma size CB+ neurons appear to be morphologically intermediate between PV+ and CR+ interneurons. While the size of CB+ neurons is similar to CR+ neurons, their dendritic topology is more similar to PV+ neurons. CB+ is also co-expressed by both CR+ and PV+ neurons, but the extent of CB co-localization with PV is greater. This is interesting in light of the fact that both PV+ and CB+ neurons originate from an overlapping embryonic pool during development (Fogarty et al., 2007). In contrast, the highly divergent morphology and CaBP expression of CR+ and PV+ neurons highlight their distinct embryonic origin (Fogarty et al., 2007; Gonchar et al., 2007; Ma et al., 2013; Radonjic et al., 2014).

### Species differences in single cell morphology of CB+ and PV+, but not CR+, interneurons

Species also had significant main effects on somato-dendritic parameters, primarily dependent on CaBP cell type. Overall, our multivariate data analyses revealed that compared to CR+ neurons, CB+ and PV+ neurons exhibited significant between-species differences in somato- dendritic morphology. Specifically, CB+ and PV+, but not CR+ neurons, exhibited a higher number of primary dendrites but smaller vertical extents and polarization in mouse V1 and FC compared to monkey V1 and LPFC. Our dimensional reduction and clustering analyses revealed that the subpopulation of CB+ and PV+ neurons from mouse and monkey were separated into subclusters. We found that monkey PV+ neurons had slightly larger somata than mouse PV+ neurons in both visual and frontal cortices. (Kooijmans et al., 2020) also found very subtle between-species differences in interneuron morphology in V1, with PV+ interneuron cell bodies slightly larger but CB neurons slightly smaller in monkey than in mouse. This scaling effect is small compared to the significant scaling we and others have observed with L3 pyramidal neurons in frontal cortices of the monkey versus mouse(e.g. Ballesteros-Yanez et al., 2007; Elston et al., 2011; Elston & DeFelipe, 2002; Gilman et al., 2017) (reviews:Elston et al., 2011; Luebke, 2017; Wittenberg, 2012). Further, we have shown that this species dependent scaling of L3 pyramidal neurons is region-specific, with neurons in LPFC monkey, but not monkey V1 being larger and more complex than in mouse FC and V1 (Gilman et al., 2017; Luebke, 2017).

In contrast to CB+ and PV+ neurons, CR+ neurons in L2-3 did not exhibit significant between-species differences in somato-dendritic morphology, and did not show subclustering of subpopulations based on species or area. These data are consistent with some previous studies which also reported no significant between-area and between-species differences in the size of CR+ neurons (Gabbott & Bacon, 1996a; Gabbott et al., 1997; Kooijmans et al., 2020). However, as shown here and in previous studies, there is a greater density and proportion of CR+ neurons in monkey compared to mouse. These data suggest that while CR+ neurons appear to be similar in morphology between species, this neuron subpopulation is differently distributed and expanded in monkeys (Dzaja et al., 2014; Hladnik et al., 2014). Interestingly, while there seems to be no dendritic scaling of CR+ neuron in L2-3 between species, the thickness of L2-3 differ across areas and species. Thus, CR+ neurons of similar dendritic vertical extent span more layers in mouse compared to monkey areas, especially when comparing frontal cortices between the two species. Indeed, we found a greater portion of CR+ neurons that spanned 2 or more layers in mouse compared to monkey areas.

Compared to species and CaBP cell type, somato-dendritic morphological features were not significantly dependent on cortical area. Of all somato-dendritic features, a significant interactive effect of area and CaBP cell type was found only for dendritic horizontal extent. Specifically CB+ and PV+, but not CR+ neurons, exhibited greater horizontal extents in frontal than in visual cortices in both species. This is interesting in light of the fact that a marked areal difference was also found with regard to the density of CB+ and PV+ inhibitory neurons in both mouse and monkey. Specifically, both CB+ and PV+ interneurons were significantly greater in density in visual compared to frontal areas. This data is consistent with previous studies showing systematic graded differences in CB+ and PV+ neuronal density across cytoarchitectonically distinct areas in monkey visual and prefrontal cortices (Dombrowski et al., 2001; Gabbott & Bacon, 1996b; John et al., 2022; Kondo et al., 1994). Interestingly, the 2x higher neuronal density in V1 compared to LPFC in monkey, is mainly due to differences in pyramidal neuron densities, which appear to vary across areas to a greater degree compared to the total interneuron population (Barbas & Pandya, 1989; Dombrowski et al., 2001; Hilgetag et al., 2016; Hsu et al., 2017; John et al., 2022; Kondo et al., 1999; O’Kusky & Colonnier, 1982). Thus, similar to dendritic scaling differences (Gilman et al., 2017; Luebke, 2017), between-area differences in interneuron densities appear to be less pronounced compared to pyramidal neuron populations in monkey V1 and LPFC.

### Implications for Species and Areal Specialization of Cortical Circuitry

The implications of major excitatory neuron specializations together with conserved interneuron properties should be explored in future studies. Our previous study at the synaptic level showed that while there is no significant species-differences in the density of inhibitory synapses, mouse visual and frontal cortices have ∼2x the density of excitatory synapses in the neuropil compared to monkey V1 and LPFC (Hsu et al., 2017). These data suggest a greater overall excitatory:inhibitory (E:I) ratio in mouse compared to monkey neuropil, where the effect of inhibition maybe stronger. Further, between-area differences in dendritic arbor size and spine density of single neurons were found to be specific to monkey cortex, implying that at the single-cell level this E:I balance differs between monkey cortical areas. Our previous studies show that in monkey, L3 V1 pyramidal neurons are specialized by having lower spine density and smaller arbors, while in LPFC pyramidal neurons have greater spine densities and larger arbors (Amatrudo et al., 2012; Gilman et al., 2017), which likely impact how inhibitory inputs are integrated. Our data here suggest that by contrast to pyramidal neurons, somato-dendritic scaling and morphology of individual interneurons likely plays a minor role in interneuron diversity across areas and species. Cortical regional and species diversity in inhibitory function is thus likely to be derived mainly from differential laminar distribution of distinct interneuron subclasses and their specific axonal connectivity profiles, which we have not examined in this study. Indeed, our previous work comparing two distinct PFC areas in monkeys showed that medial PFC had a significantly greater inhibitory tone compared to LPFC, due to different perisomatic inhibitory synapses (Medalla et al., 2017). Computational modeling incorporating these empirically-derived properties show that this difference in E:I synaptic balance can lead to area-specific cortical dynamics (Medalla et al., 2017). The distribution of inhibitory inputs along somato-dendritic compartments of pyramidal neurons in mouse versus monkey frontal and visual cortices remain largely unknown and will likely have an impact on circuit activity required to perform species and cortical area specific functions.

The data presented here adds to the growing literature on how nuanced differences in the population organization and individual properties of constituent neurons may underlie specializations in cortical regions responsible for species and area-specific functional capacities. Taken together with work showing differences in the cytoarchitecture of different regions and in the density and distribution of pyramidal neurons between species (Barbas & Pandya, 1989; Hsu et al., 2017; Keller et al., 2018; O’Kusky & Colonnier, 1982; Petrides et al., 2012; Rockland & Pandya, 1979; Van De Werd et al., 2010; Wang & Burkhalter, 2007) (review: Pandya, 1985; Yeterian et al., 2012), a clearer picture is emerging about which are the key specializations of principal cells -particularly pyramidal cells- versus interneurons that may underlie areal functional specialization. While we have shown that the density and distribution of CaBP+ interneurons does differ between areas and species and may be important, we can also rule out that major differences in somato-dendritic features of interneurons underlie species and area differences. Indeed, the somato-dendritic morphology of interneurons did not substantially scale across the neuroaxis in the monkey as pyramidal neurons do- differences were much more dependent on CaBP type than on area or species, by contrast to what has been shown for L3 pyramidal cells (Gilman et al., 2017; Luebke, 2017). In contrast to pyramidal cells which differ widely in distribution and morphology across areas and species, the distribution and morphology of interneurons -while differing in many details- appears to be far more conserved across areas and species.

## Acknowledgements

This work was supported by NIH/NIA R01-AG071230; R01-AG059028 (JL), and NIH/NIMH R01-MH116008 (MM). The authors are grateful for the continuous collaborative support of our colleague Dr. Doug Rosene.

## References

1. Amatrudo, J. M., Weaver, C. M., Crimins, J. L., Hof, P. R., Rosene, D. L., & Luebke, J. I. (2012). Influence of highly distinctive structural properties on the excitability of pyramidal neurons in monkey visual and prefrontal cortices. J Neurosci, 32(40), 13644–13660. doi:10.1523/JNEUROSCI.2581-12.2012

2. Anastasiades, P. G., Boada, C., & Carter, A. G. (2019). Cell-Type-Specific D1 Dopamine Receptor Modulation of Projection Neurons and Interneurons in the Prefrontal Cortex. Cereb Cortex, 29(7), 3224–3242. doi:10.1093/cercor/bhy299

3. Ballesteros-Yanez, I., Ambrosio, E., Benavides-Piccione, R., Perez, J., Torres, I., Miguens, M., . . . DeFelipe, J. (2007). The effects of morphine self-administration on cortical pyramidal cell structure in addiction-prone lewis rats. Cereb Cortex, 17(1), 238–249. doi:10.1093/cercor/bhj142

4. Barbas, H., Medalla, M., Alade, O., Suski, J., Zikopoulos, B., & Lera, P. (2005). Relationship of prefrontal connections to inhibitory systems in superior temporal areas in the rhesus monkey. Cereb Cortex, 15(9), 1356–1370. doi:10.1093/cercor/bhi018

5. Barbas, H., & Pandya, D. N. (1989). Architecture and intrinsic connections of the prefrontal cortex in the rhesus monkey. J Comp Neurol, 286(3), 353–375. doi:10.1002/cne.902860306

6. Barinka, F., Salaj, M., Rybar, J., Krajcovicova, E., Kubova, H., & Druga, R. (2012). Calretinin, parvalbumin and calbindin immunoreactive interneurons in perirhinal cortex and temporal area Te3V of the rat brain: qualitative and quantitative analyses. Brain Res, 1436, 68–80. doi:10.1016/j.brainres.2011.12.014

7. Beaulieu, C. (1993). Numerical data on neocortical neurons in adult rat, with special reference to the GABA population. Brain Res, 609(1-2), 284–292. doi:10.1016/0006-8993(93)90884-p

8. Buzsaki, G., Geisler, C., Henze, D. A., & Wang, X. J. (2004). Interneuron Diversity series: Circuit complexity and axon wiring economy of cortical interneurons. Trends Neurosci, 27(4), 186–193. doi:10.1016/j.tins.2004.02.007

9. Cauli, B., Audinat, E., Lambolez, B., Angulo, M. C., Ropert, N., Tsuzuki, K., . . . Rossier, J. (1997). Molecular and physiological diversity of cortical nonpyramidal cells. J Neurosci, 17(10), 3894–3906. doi:10.1523/JNEUROSCI.17-10-03894.1997

10. Cauli, B., Zhou, X., Tricoire, L., Toussay, X., & Staiger, J. F. (2014). Revisiting enigmatic cortical calretinin-expressing interneurons. Front Neuroanat, 8, 52. doi:10.3389/fnana.2014.00052

11. Conde, F., Lund, J. S., Jacobowitz, D. M., Baimbridge, K. G., & Lewis, D. A. (1994). Local circuit neurons immunoreactive for calretinin, calbindin D-28k or parvalbumin in monkey prefrontal cortex: distribution and morphology. J Comp Neurol, 341(1), 95–116. doi:10.1002/cne.903410109

12. DeFelipe, J. (1997). Types of neurons, synaptic connections and chemical characteristics of cells immunoreactive for calbindin-D28K, parvalbumin and calretinin in the neocortex. J Chem Neuroanat, 14(1), 1–19.

13. DeFelipe, J. (2002). Cortical interneurons: from Cajal to 2001. Progress in Brain Research, 136, 215–238.

14. Defelipe, J. (2011). The evolution of the brain, the human nature of cortical circuits, and intellectual creativity. Front Neuroanat, 5, 29. doi:10.3389/fnana.2011.00029

15. DeFelipe, J., Gonzalez-Albo, M. C., del Rio, M. R., & Elston, G. N. (1999). Distribution and patterns of connectivity of interneurons containing calbindin, calretinin, and parvalbumin in visual areas of the occipital and temporal lobes of the macaque monkey. Journal of Comparative Neurology, 412, 515–526.

16. DeFelipe, J., Hendry, S. H., Hashikawa, T., Molinari, M., & Jones, E. G. (1990). A microcolumnar structure of monkey cerebral cortex revealed by immunocytochemical studies of double bouquet cell axons. Neuroscience, 37, 655–673.

17. DeFelipe, J., Lopez-Cruz, P. L., Benavides-Piccione, R., Bielza, C., Larranaga, P., Anderson, S., . . . Ascoli, G. A. (2013). New insights into the classification and nomenclature of cortical GABAergic interneurons. Nat Rev Neurosci, 14(3), 202–216. doi:10.1038/nrn3444

18. Desgent, S., Boire, D., & Ptito, M. (2005). Distribution of calcium binding proteins in visual and auditory cortices of hamsters. Exp Brain Res, 163(2), 159–172. doi:10.1007/s00221-004-2151-3

19. Disney, A. A., & Aoki, C. (2008). Muscarinic acetylcholine receptors in macaque V1 are most frequently expressed by parvalbumin-immunoreactive neurons. J Comp Neurol, 507(5), 1748–1762. doi:10.1002/cne.21616

20. Disney, A. A., & Reynolds, J. H. (2014). Expression of m1-type muscarinic acetylcholine receptors by parvalbumin-immunoreactive neurons in the primary visual cortex: a comparative study of rat, guinea pig, ferret, macaque, and human. J Comp Neurol, 522(5), 986–1003. doi:10.1002/cne.23456

21. Dombrowski, S. M., Hilgetag, C. C., & Barbas, H. (2001). Quantitative architecture distinguishes prefrontal cortical systems in the rhesus monkey. Cerebral Cortex, 11, 975–988.

22. Dzaja, D., Hladnik, A., Bicanic, I., Bakovic, M., & Petanjek, Z. (2014). Neocortical calretinin neurons in primates: increase in proportion and microcircuitry structure. Front Neuroanat, 8, 103. doi:10.3389/fnana.2014.00103

23. Elston, G. N., Benavides-Piccione, R., Elston, A., Manger, P. R., & Defelipe, J. (2011). Pyramidal cells in prefrontal cortex of primates: marked differences in neuronal structure among species. Front Neuroanat, 5, 2. doi:10.3389/fnana.2011.00002

24. Elston, G. N., & DeFelipe, J. (2002). Spine distribution in cortical pyramidal cells: a common organizational principle across species. Progress in Brain Research, 136, 109–133.

25. Fogarty, M., Grist, M., Gelman, D., Marin, O., Pachnis, V., & Kessaris, N. (2007). Spatial genetic patterning of the embryonic neuroepithelium generates GABAergic interneuron diversity in the adult cortex. J Neurosci, 27(41), 10935–10946. doi:10.1523/JNEUROSCI.1629-07.2007

26. Gabbott, P. L., & Bacon, S. J. (1996a). Local circuit neurons in the medial prefrontal cortex (areas 24a,b,c, 25 and 32) in the monkey: I. Cell morphology and morphometrics. J Comp Neurol, 364(4), 567–608. doi:10.1002/(SICI)1096-9861(19960122)364:4<567::AID- CNE1>3.0.CO;2-1

27. Gabbott, P. L., & Bacon, S. J. (1996b). Local circuit neurons in the medial prefrontal cortex (areas 24a,b,c, 25 and 32) in the monkey: II. Quantitative areal and laminar distributions. J Comp Neurol, 364(4), 609–636. doi:10.1002/(SICI)1096-9861(19960122)364:4<609::AID- CNE2>3.0.CO;2-7

28. Gabbott, P. L., Dickie, B. G., Vaid, R. R., Headlam, A. J., & Bacon, S. J. (1997). Local-circuit neurones in the medial prefrontal cortex (areas 25, 32 and 24b) in the rat: morphology and quantitative distribution. J Comp Neurol, 377(4), 465–499. doi:10.1002/(sici)1096- 9861(19970127)377:4<465::aid-cne1>3.0.co;2-0

29. Gilman, J. P., Medalla, M., & Luebke, J. I. (2017). Area-Specific Features of Pyramidal Neurons-a Comparative Study in Mouse and Rhesus Monkey. Cereb Cortex, 27(3), 2078–2094. doi:10.1093/cercor/bhw062

30. Glezer, II, Hof, P. R., & Morgane, P. J. (1998). Comparative analysis of calcium-binding protein-immunoreactive neuronal populations in the auditory and visual systems of the bottlenose dolphin (Tursiops truncatus) and the macaque monkey (Macaca fascicularis). J Chem Neuroanat, 15(4), 203–237. doi:10.1016/s0891-0618(98)00022-2

31. Gonchar, Y., & Burkhalter, A. (1997). Three distinct families of GABAergic neurons in rat visual cortex. Cereb Cortex, 7(4), 347–358. doi:10.1093/cercor/7.4.347

32. Gonchar, Y., Wang, Q., & Burkhalter, A. (2007). Multiple distinct subtypes of GABAergic neurons in mouse visual cortex identified by triple immunostaining. Front Neuroanat, 1, 3. doi:10.3389/neuro.05.003.2007

33. Guillamon-Vivancos, T., Tyler, W. A., Medalla, M., Chang, W. W., Okamoto, M., Haydar, T. F., & Luebke, J. I. (2019). Distinct Neocortical Progenitor Lineages Fine-tune Neuronal Diversity in a Layer-specific Manner. Cereb Cortex, 29(3), 1121–1138. doi:10.1093/cercor/bhy019

34. Hilgetag, C. C., Medalla, M., Beul, S. F., & Barbas, H. (2016). The primate connectome in context: Principles of connections of the cortical visual system. Neuroimage, 134, 685–702. doi:10.1016/j.neuroimage.2016.04.017

35. Hladnik, A., Dzaja, D., Darmopil, S., Jovanov-Milosevic, N., & Petanjek, Z. (2014). Spatio-temporal extension in site of origin for cortical calretinin neurons in primates. Front Neuroanat, 8, 50. doi:10.3389/fnana.2014.00050

36. Hof, P. R., Glezer, II, Conde, F., Flagg, R. A., Rubin, M. B., Nimchinsky, E. A., & Vogt Weisenhorn, D. M. (1999). Cellular distribution of the calcium-binding proteins parvalbumin, calbindin, and calretinin in the neocortex of mammals: phylogenetic and developmental patterns. J Chem Neuroanat, 16(2), 77–116. doi:10.1016/s0891-0618(98)00065-9

37. Hsu, A., Luebke, J. I., & Medalla, M. (2017). Comparative ultrastructural features of excitatory synapses in the visual and frontal cortices of the adult mouse and monkey. J. Comp. Neurol., 525(9), 2175–2191. doi:10.1002/cne.24196

38. Isaacson, J. S., & Scanziani, M. (2011). How inhibition shapes cortical activity. Neuron, 72(2), 231–243. doi:10.1016/j.neuron.2011.09.027

39. John, Y. J., Zikopoulos, B., Garcia-Cabezas, M. A., & Barbas, H. (2022). The cortical spectrum: A robust structural continuum in primate cerebral cortex revealed by histological staining and magnetic resonance imaging. Front Neuroanat, 16, 897237. doi:10.3389/fnana.2022.897237

40. Jones, E. G. H., S.H. (1986). Co-localization of GABA and neuropeptides in neocortical neurons. Trends Neurosci, 9, 71–76. doi:https://doi.org/10.1016/0166-2236(86)90026-3

41. Kawaguchi, Y., & Kubota, Y. (1997). GABAergic cell subtypes and their synaptic connections in rat frontal cortex. Cerebral Cortex, 7, 476–486.

42. Keller, D., Ero, C., & Markram, H. (2018). Cell Densities in the Mouse Brain: A Systematic Review. Front Neuroanat, 12, 83. doi:10.3389/fnana.2018.00083

43. Kondo, H., Hashikawa, T., Tanaka, K., & Jones, E. G. (1994). Neurochemical gradient along the monkey occipito-temporal cortical pathway. NeuroReport, 5(5), 613–616.

44. Kondo, H., Tanaka, K., Hashikawa, T., & Jones, E. G. (1999). Neurochemical gradients along monkey sensory cortical pathways: calbindin-immunoreactive pyramidal neurons in layers II and III. Eur.J Neurosci., 11(12), 4197–4203.

45. Kooijmans, R. N., Self, M. W., Wouterlood, F. G., Belien, J. A., & Roelfsema, P. R. (2014). Inhibitory interneuron classes express complementary AMPA-receptor patterns in macaque primary visual cortex. J Neurosci, 34(18), 6303–6315. doi:10.1523/JNEUROSCI.3188-13.2014

46. Kooijmans, R. N., Sierhuis, W., Self, M. W., & Roelfsema, P. R. (2020). A Quantitative Comparison of Inhibitory Interneuron Size and Distribution between Mouse and Macaque V1, Using Calcium-Binding Proteins. Cereb Cortex Commun, 1(1), tgaa068. doi:10.1093/texcom/tgaa068

47. Kubota, Y. (2014). Untangling GABAergic wiring in the cortical microcircuit. Curr Opin Neurobiol, 26, 7–14. doi:10.1016/j.conb.2013.10.003

48. Kubota, Y., Karube, F., Nomura, M., & Kawaguchi, Y. (2016). The Diversity of Cortical Inhibitory Synapses. Front Neural Circuits, 10, 27. doi:10.3389/fncir.2016.00027

49. LeBlang, C. J., Medalla, M., Nicoletti, N. W., Hays, E. C., Zhao, J., Shattuck, J., . . . Luebke, J. I. (2020). Reduction of the RNA Binding Protein TIA1 Exacerbates Neuroinflammation in Tauopathy. Front Neurosci, 14, 285. doi:10.3389/fnins.2020.00285

50. Lourenco, J., Koukouli, F., & Bacci, A. (2020). Synaptic inhibition in the neocortex: Orchestration and computation through canonical circuits and variations on the theme. Cortex, 132, 258–280. doi:10.1016/j.cortex.2020.08.015

51. Luebke, J. I. (2017). Pyramidal Neurons Are Not Generalizable Building Blocks of Cortical Networks. Front. Neuroanat., 11, 11. doi:10.3389/fnana.2017.00011

52. Ma, T., Wang, C., Wang, L., Zhou, X., Tian, M., Zhang, Q., . . . Yang, Z. (2013). Subcortical origins of human and monkey neocortical interneurons. Nat Neurosci, 16(11), 1588–1597. doi:10.1038/nn.3536

53. Markram, H., Toledo-Rodriguez, M., Wang, Y., Gupta, A., Silberberg, G., & Wu, C. (2004). Interneurons of the neocortical inhibitory system. Nat Rev Neurosci 5(10), 793–807.

54. Medalla, M., & Barbas, H. (2010). Anterior cingulate synapses in prefrontal areas 10 and 46 suggest differential influence in cognitive control. J Neurosci, 30(48), 16068–16081. doi:10.1523/JNEUROSCI.1773-10.2010

55. Medalla, M., Chang, W., Calderazzo, S. M., Go, V., Tsolias, A., Goodliffe, J. W., . . . Moore, T. L. (2020). Treatment with Mesenchymal-Derived Extracellular Vesicles Reduces Injury-Related Pathology in Pyramidal Neurons of Monkey Perilesional Ventral Premotor Cortex. J Neurosci, 40(17), 3385–3407. doi:10.1523/JNEUROSCI.2226-19.2020

56. Medalla, M., Chang, W., Ibanez, S., Guillamon-Vivancos, T., Nittmann, M., Kapitonava, A., . . . Luebke, J. I. (2021). Layer-specific pyramidal neuron properties underlie diverse anterior cingulate cortical motor and limbic networks. Cereb Cortex. doi:10.1093/cercor/bhab347

57. Medalla, M., Gilman, J. P., Wang, J. Y., & Luebke, J. I. (2017). Strength and Diversity of Inhibitory Signaling Differentiates Primate Anterior Cingulate from Lateral Prefrontal Cortex. J Neurosci, 37(18), 4717–4734. doi:10.1523/JNEUROSCI.3757-16.2017

58. Meinecke, D. L., & Peters, A. (1987). GABA immunoreactive neurons in rat visual cortex. J Comp Neurol, 261(3), 388–404. doi:10.1002/cne.902610305

59. Meskenaite, V. (1997). Calretinin-immunoreactive local circuit neurons in area 17 of the cynomolgus monkey, Macaca fascicularis. J Comp Neurol, 379(1), 113–132.

60. Monyer, H., & Markram, H. (2004). Interneuron Diversity series: Molecular and genetic tools to study GABAergic interneuron diversity and function. Trends Neurosci, 27(2), 90–97. doi:10.1016/j.tins.2003.12.008

61. O’Kusky, J., & Colonnier, M. (1982). A laminar analysis of the number of neurons, glia, and synapses in the adult cortex (area 17) of adult macaque monkeys. J Comp Neurol, 210(3), 278–290. doi:10.1002/cne.902100307

62. Pandya, D. N. Y., E. H. (1985). Architecture and Connections of Cortical Association Areas. In: Peters, A., Jones, E.G. (eds) Association and Auditory Cortices. In A. J. Peters, E.G. (Ed.), Cerebral Cortex (Vol. 4). Boston, MA: Springer.

63. Petrides, M., Tomaiuolo, F., Yeterian, E. H., & Pandya, D. N. (2012). The prefrontal cortex: comparative architectonic organization in the human and the macaque monkey brains. Cortex, 48(1), 46–57. doi:10.1016/j.cortex.2011.07.002

64. Povysheva, N. V., Zaitsev, A. V., Rotaru, D. C., Gonzalez-Burgos, G., Lewis, D. A., & Krimer, L. S. (2008). Parvalbumin-positive basket interneurons in monkey and rat prefrontal cortex. J Neurophysiol, 100(4), 2348–2360. doi:10.1152/jn.90396.2008

65. Radonjic, N. V., Ortega, J. A., Memi, F., Dionne, K., Jakovcevski, I., & Zecevic, N. (2014). The complexity of the calretinin-expressing progenitors in the human cerebral cortex. Front Neuroanat, 8, 82. doi:10.3389/fnana.2014.00082

66. Rockland, K. S., & Pandya, D. N. (1979). Laminar origins and terminations of cortical connections of the occipital lobe in the rhesus monkey. Brain Res, 179(1), 3–20. doi:10.1016/0006-8993(79)90485-2

67. Sherwood, C. C., Raghanti, M. A., Stimpson, C. D., Bonar, C. J., de Sousa, A. A., Preuss, T. M., & Hof, P. R. (2007). Scaling of inhibitory interneurons in areas v1 and v2 of anthropoid primates as revealed by calcium-binding protein immunohistochemistry. Brain Behav Evol, 69(3), 176–195. doi:10.1159/000096986

68. Tremblay, R., Lee, S., & Rudy, B. (2016). GABAergic Interneurons in the Neocortex: From Cellular Properties to Circuits. Neuron, 91(2), 260–292. doi:10.1016/j.neuron.2016.06.033

69. Tsolias, A., & Medalla, M. (2022). Muscarinic Acetylcholine Receptor Localization on Distinct Excitatory and Inhibitory Neurons Within the ACC and LPFC of the Rhesus Monkey. Front Neural Circuits, 15, 795325. doi:10.3389/fncir.2021.795325

70. Uylings, H. B., Groenewegen, H. J., & Kolb, B. (2003). Do rats have a prefrontal cortex? Behav Brain Res, 146(1-2), 3–17. doi:10.1016/j.bbr.2003.09.028

71. van Brederode, J. F., Mulligan, K. A., & Hendrickson, A. E. (1990). Calcium-binding proteins as markers for subpopulations of GABAergic neurons in monkey striate cortex. Journal of Comparative Neurology, 298, 1–22.

72. Van De Werd, H. J., Rajkowska, G., Evers, P., & Uylings, H. B. (2010). Cytoarchitectonic and chemoarchitectonic characterization of the prefrontal cortical areas in the mouse. Brain Struct Funct, 214(4), 339–353. doi:10.1007/s00429-010-0247-z

73. Wang, Q., & Burkhalter, A. (2007). Area map of mouse visual cortex. J Comp Neurol, 502(3), 339–357. doi:10.1002/cne.21286

74. Winsky, L., & Kuznicki, J. (1996). Antibody recognition of calcium-binding proteins depends on their calcium-binding status. J Neurochem, 66(2), 764–771. doi:10.1046/j.1471-4159.1996.66020764.x

75. Wittenberg, G. M. W., S. S. H. . (2012). Evolution and scaling of dendrites. . In G. S. Stuart, N.; Hausser, M. (Ed.), Dendrites (2nd ed.): Oxford University Press.

76. Xu, X., Olivas, N. D., Ikrar, T., Peng, T., Holmes, T. C., Nie, Q., & Shi, Y. (2016). Primary visual cortex shows laminar-specific and balanced circuit organization of excitatory and inhibitory synaptic connectivity. J Physiol, 594(7), 1891–1910. doi:10.1113/JP271891

77. Yeterian, E. H., Pandya, D. N., Tomaiuolo, F., & Petrides, M. (2012). The cortical connectivity of the prefrontal cortex in the monkey brain. Cortex, 48(1), 58–81. doi:10.1016/j.cortex.2011.03.004

78. Yuste, R. (2005). Origin and classification of neocortical interneurons. Neuron, 48(4), 524–527. doi:10.1016/j.neuron.2005.11.012

79. Zaitsev, A. V., Gonzalez-Burgos, G., Povysheva, N. V., Kroner, S., Lewis, D. A., & Krimer, L. S. (2005). Localization of calcium-binding proteins in physiologically and morphologically characterized interneurons of monkey dorsolateral prefrontal cortex. Cereb Cortex, 15(8), 1178–1186. doi:10.1093/cercor/bhh218

80. Zaitsev, A. V., Povysheva, N. V., Gonzalez-Burgos, G., Rotaru, D., Fish, K. N., Krimer, L. S., & Lewis, D. A. (2009). Interneuron diversity in layers 2-3 of monkey prefrontal cortex. Cereb Cortex, 19(7), 1597–1615. doi:10.1093/cercor/bhn198

