## Supplemental Material for "Comparative Features of Calretinin, Calbindin and Parvalbumin Expressing Interneurons in Mouse and Monkey Primary Visual and Frontal Cortices"

### Supporting Information

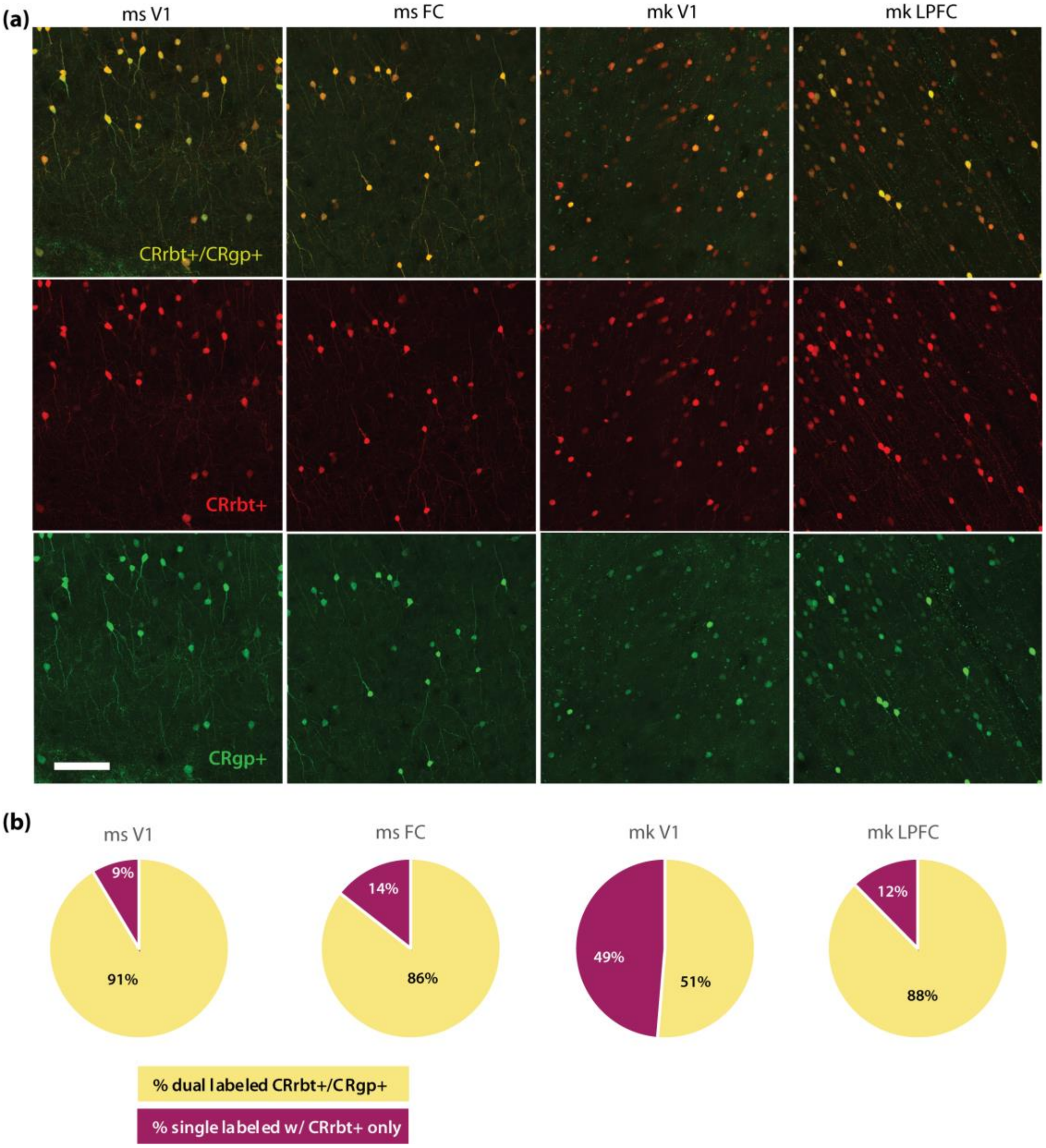


**Supplementary Figure S1. Variability of calretinin labeling based on antibody species.** A) Confocal images showing dual labeling of CR+ neurons using rabbit anti-CR (CRrbt, Swant, 7697; RRID AB_2721226) and guinea pig anti-CR (CRgp, Swant CRgp7) primary antibodies in one monkey and one mouse case. Scale bar: 100µm. B) Pie charts showing the proportion of cells dual labeled with the two CR+ antibodies (CRrbt+/CRgp+), and cells single labeled with only the CR-rabbit antibody.

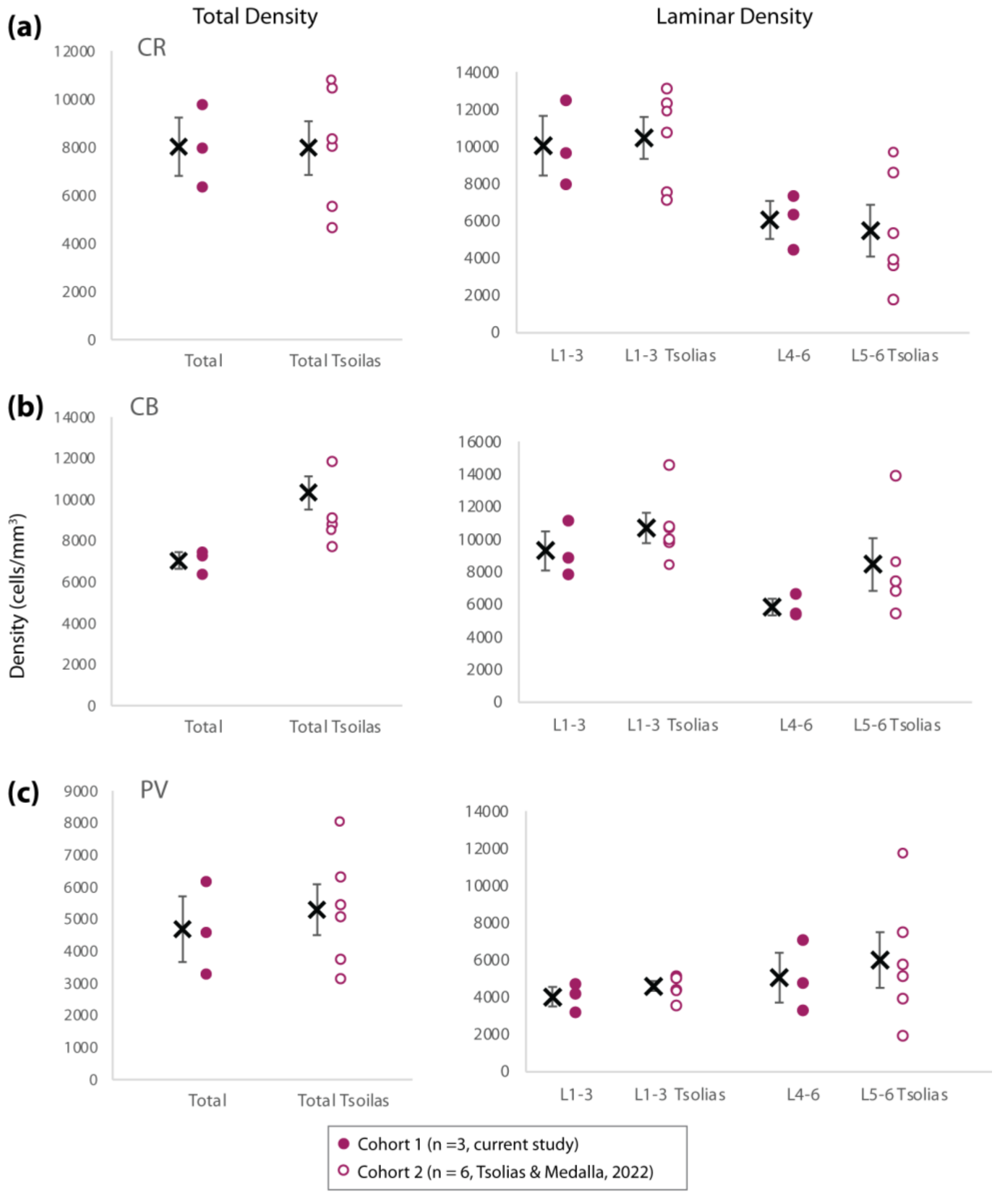


**Supplementary Figure S2. Comparison of CR+ CB+ PV+ neuronal density quantification in LPFC area 46 from two independent cohorts of monkeys by independent investigators.** Total density and density in upper (L1-3) and deep (L4-6 or L5-6) layers of LPFC area 46 of A) CR+ neurons, B) CB+ neurons and, C) PV+ neurons. These data show that quantification from the cohort (n =3) in the present study is consistent with quantification in a second larger (n = 6) cohort analyzed by a different investigator as part of our previous study (Tsolias and Medalla, 2022). We do note that there is a slight difference in deep layer quantification with the cohort (cohort 1) in the present study including cell density counts in layer 4, while the previous cohort (cohort 2) quantified cells within layers 5-6. Nonetheless statistical comparison by paired t-test, shows no significant differences between the cohorts with regards to density of CR+, CB+ and PV+ in LPFC area 46.

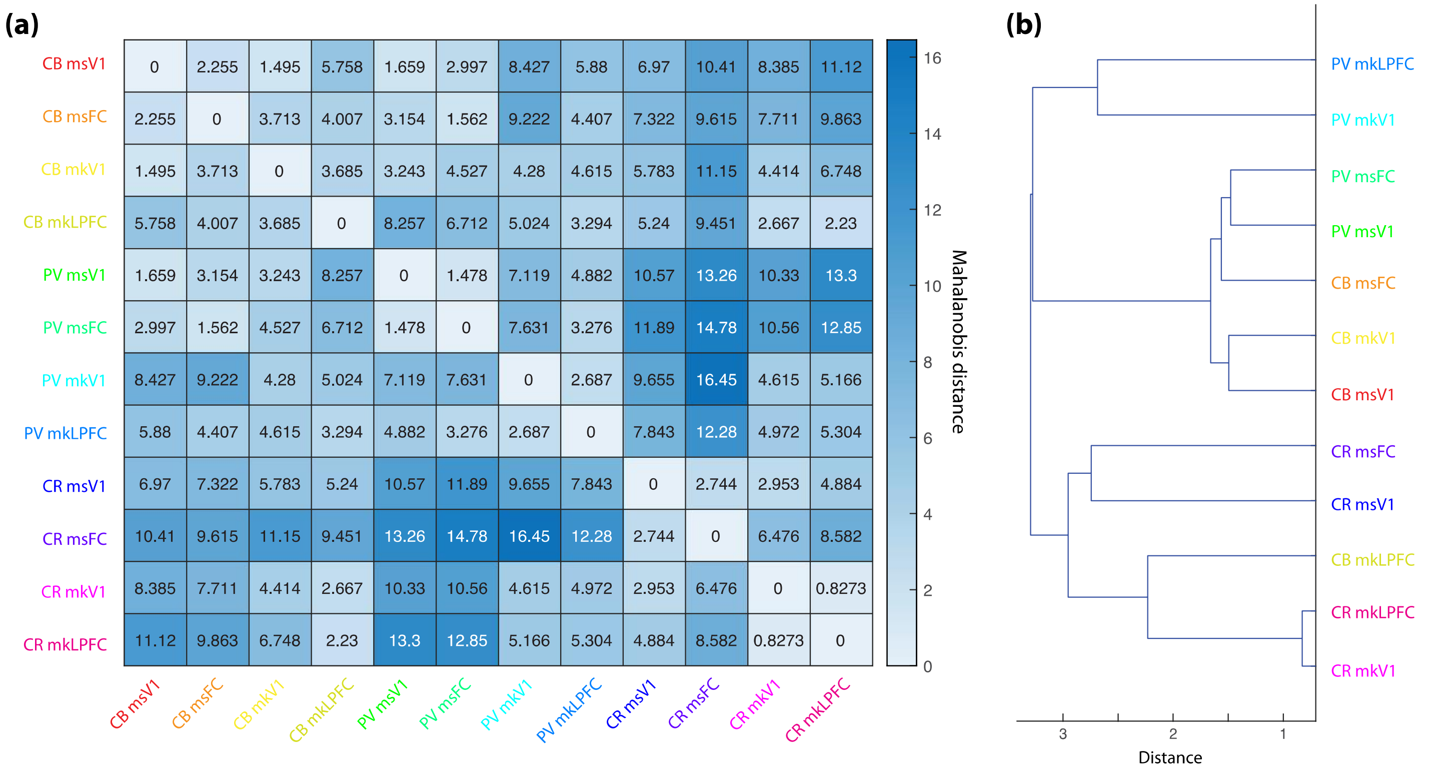


**Supplementary Figure S3. Distance matrix and clustering of group means based on MANOVA.** A) Mahalanobis distance matrix of group means derived from MANOVA of multivariate somato-dendritic properties, showing relative (dis)similarities between interneuron groups by CaBP expression (CB, PV, CR), species (mouse, ms vs monkey, mk) and cortical area (visual V1 vs frontal FC, LPFC). B) Dendrogram showing clustering of group means derived by applying the single linkage method to the matrix of Mahalanobis distances between group means.

### Table S1: 3-way ANOVA main effects and Fischer's LSD post-hoc test p-values for morphological parameters.

| | Soma Volume | # of Primary Dendrites | # Layers Spanned | 2D Convex Hull Area | Horizontal Extent | Vertical Extent | Polarity [X] | Polarity [Y] |
|---|---|---|---|---|---|---|---|---|
| ***Species*** | $3.00 \times 10^{-8}$ | $1.15 \times 10^{-15}$ | $3.76 \times 10^{-8}$ | 0.072 | $1.56 \times 10^{-5}$ | 0.013 | 0.086 | $1.18 \times 10^{-8}$ |
| ***Area*** | 0.139 | 0.629 | 0.438 | $8.04 \times 10^{-7}$ | $3.63 \times 10^{-9}$ | $1.32 \times 10^{-11}$ | 0.750 | 0.091 |
| ***CaBP*** | $3.05 \times 10^{-18}$ | $4.76 \times 10^{-48}$ | $6.98 \times 10^{-21}$ | 0.003 | $1.66 \times 10^{-5}$ | $1.83 \times 10^{-15}$ | 0.143 | 0.019 |
| ***Species*Area*** | 0.418 | 0.671 | 0.499 | 0.349 | 0.080 | 0.131 | 0.860 | 0.463 |
| ***Spec*CaBP*** | 0.022 | $2.39 \times 10^{-4}$ | $1.41 \times 10^{-7}$ | 0.010 | 0.408 | 0.020 | 0.065 | 0.773 |
| ***Area*CaBP*** | 0.548 | 0.005 | 0.263 | 0.007 | 0.002 | 0.392 | 0.126 | 0.796 |
| **Between-species** | | | | | | | | |
| **V1 ms vs mk** | | | | | | | | |
| CR | 0.001 | 0.315 | $1.97 \times 10^{-9}$ | 0.007 | 0.641 | 0.184 | 0.008 | 0.060 |
| CB | 0.150 | $5.54 \times 10^{-6}$ | 0.505 | 0.061 | 0.036 | 0.404 | 0.431 | 0.007 |
| PV | $5.03 \times 10^{-6}$ | $1.57 \times 10^{-9}$ | 0.921 | 0.468 | 0.191 | 0.039 | 0.661 | 0.004 |
| **FC ms vs mk** | | | | | | | | |
| CR | 0.006 | 0.141 | $4.43 \times 10^{-12}$ | 0.052 | 0.028 | 0.945 | 0.023 | 0.007 |
| CB | 0.459 | $1.30 \times 10^{-8}$ | 0.124 | 0.272 | $1.28 \times 10^{-5}$ | 0.007 | 0.766 | $9.38 \times 10^{-5}$ |
| PV | $1.15 \times 10^{-4}$ | $5.55 \times 10^{-11}$ | 0.428 | 0.090 | 0.002 | $2.86 \times 10^{-4}$ | 0.319 | $2.36 \times 10^{-4}$ |
| **Between-area** | | | | | | | | |
| **ms V1 vs FC** | | | | | | | | |
| CR | 0.639 | 0.580 | 0.096 | 0.926 | 0.195 | 0.021 | 0.930 | 0.269 |
| CB | 0.063 | 0.027 | 0.820 | 0.015 | $2.67 \times 10^{-6}$ | $2.62 \times 10^{-4}$ | 0.093 | 0.133 |
| PV | 0.284 | 0.141 | 0.549 | $2.11 \times 10^{-4}$ | $7.65 \times 10^{-8}$ | 0.040 | 0.739 | 0.030 |
| **mk V1 vs LPFC** | | | | | | | | |
| CR | 0.768 | 0.869 | 0.280 | 0.418 | 0.730 | $1.29 \times 10^{-4}$ | 0.541 | 0.990 |
| CB | 0.263 | 0.005 | 0.330 | $3.15 \times 10^{-4}$ | 0.001 | $4.47 \times 10^{-8}$ | 0.017 | 0.735 |
| PV | 0.758 | 0.248 | 0.947 | $5.72 \times 10^{-7}$ | $4.64 \times 10^{-5}$ | $1.31 \times 10^{-4}$ | 0.834 | 0.263 |
| **Between-CaBP cell type** | | | | | | | | |
| **CR vs CB** | | | | | | | | |
| ms V1 | 0.212 | $1.79 \times 10^{-20}$ | $7.44 \times 10^{-11}$ | 0.001 | 0.516 | $5.88 \times 10^{-8}$ | 0.180 | 0.926 |
| ms FC | 0.974 | $3.16 \times 10^{-12}$ | $7.82 \times 10^{-16}$ | 0.348 | $3.30 \times 10^{-4}$ | $1.93 \times 10^{-6}$ | 0.931 | 0.402 |
| mk V1 | 0.002 | $1.03 \times 10^{-9}$ | 0.223 | 0.029 | 0.430 | $2.55 \times 10^{-4}$ | 0.609 | 0.613 |
| mk LPFC | 0.008 | $1.23 \times 10^{-7}$ | $4.18 \times 10^{-5}$ | 0.988 | 0.004 | $1.96 \times 10^{-4}$ | 0.009 | 0.765 |
| **CR vs PV** | | | | | | | | |
| ms V1 | 0.008 | $1.89 \times 10^{-25}$ | $3.65 \times 10^{-13}$ | 0.001 | 0.261 | $1.26 \times 10^{-7}$ | 0.539 | 0.108 |
| ms FC | 0.003 | $3.70 \times 10^{-26}$ | $5.32 \times 10^{-15}$ | 0.947 | $3.82 \times 10^{-6}$ | $1.02 \times 10^{-7}$ | 0.339 | 0.057 |
| mk V1 | 0.001 | $4.36 \times 10^{-10}$ | 0.123 | 0.867 | 0.700 | 0.033 | 0.021 | 0.421 |
| mk LPFC | $9.68 \times 10^{-7}$ | $7.55 \times 10^{-18}$ | 0.001 | $4.60 \times 10^{-5}$ | $4.46 \times 10^{-7}$ | 0.002 | 0.016 | 0.133 |
| **CB vs PV** | | | | | | | | |
| ms V1 | $8.89 \times 10^{-5}$ | 0.157 | 0.399 | 0.797 | 0.631 | 0.565 | 0.462 | 0.125 |
| ms FC | 0.001 | $9.55 \times 10^{-7}$ | 0.939 | 0.286 | 0.182 | 0.385 | 0.251 | 0.222 |
| mk V1 | $5.53 \times 10^{-12}$ | 0.973 | 0.744 | 0.026 | 0.198 | 0.079 | 0.053 | 0.153 |
| mk LPFC | $4.68 \times 10^{-14}$ | $1.69 \times 10^{-5}$ | 0.563 | $1.60 \times 10^{-5}$ | 0.010 | 0.719 | 0.976 | 0.194 |

**Table S2. Coefficients of Canonical Variables from MANOVA of somato-dendritic properties of 12 groups.** Table of outcome measures used for calculating MANOVA canonical variables and corresponding coefficients for each outcome measure representing the degree of influence on separation of 12 groups of interneurons. In red are shown the largest absolute values of coefficients that had the strongest influence on each of the significant canonical variables. The eigenvalues and p values for each canonical variable are shown.

| | Coefficients (c) for Canonical Variables | | | | | | | |
|---|---|---|---|---|---|---|---|---|
| | c1 | c2 | c3 | c4 | c5 | c6 | c7 | c8 |
| **Outcome Measures:** | | | | | | | | |
| # of Layers Spanned | 0.092 | 0.962 | 0.840 | 0.260 | 0.182 | -0.174 | -0.075 | 0.420 |
| # of Primary Dendrites | 1.160 | 0.024 | 0.109 | 0.153 | 0.333 | 0.814 | 0.131 | 0.197 |
| Soma Volume(µm³) | -0.165 | -0.997 | 0.728 | 0.522 | 0.206 | -0.290 | -0.168 | 0.099 |
| Soma Area (µm²) | 0.139 | 0.196 | 0.073 | 0.179 | -1.344 | 0.477 | 0.693 | -0.396 |
| Horizontal Extent (µm) | 0.260 | 0.247 | -0.806 | 0.095 | 0.387 | -0.990 | -0.142 | 0.593 |
| Vertical Extent (µm) | -0.574 | -0.155 | -0.811 | 0.389 | 0.840 | 0.682 | -0.089 | 0.018 |
| Polarity ABS [X] | 0.111 | 0.016 | 0.124 | -0.082 | 0.348 | -0.296 | 0.915 | -0.051 |
| Polarity ABS [Y] | -0.086 | -0.223 | -0.024 | -0.452 | -0.221 | 0.157 | 0.228 | 0.873 |
| *eigenvalues* | *1.359* | *0.791* | *0.329* | *0.246* | *0.075* | *0.046* | 0.033 | 0.010 |
| *p-values* | $2.2\times10^{-112}$ | $1.3\times10^{-59}$ | $1.4\times10^{-26}$ | $3\times10^{-13}$ | *0.0005* | *0.019* | 0.110 | 0.471 |
